# GlobalFungi: Global database of fungal records from high-throughput-sequencing metabarcoding studies

**DOI:** 10.1101/2020.04.24.060384

**Authors:** Tomáš Větrovský, Daniel Morais, Petr Kohout, Clémentine Lepinay, Camelia Algora Gallardo, Sandra Awokunle Hollá, Barbara Doreen Bahnmann, Květa Bílohnědá, Vendula Brabcová, Federica D’Alò, Zander Rainier Human, Mayuko Jomura, Miroslav Kolařík, Jana Kvasničková, Salvador Lladó, Rubén López-Mondéjar, Tijana Martinović, Tereza Mašínová, Lenka Meszárošová, Lenka Michalčíková, Tereza Michalová, Sunil Mundra, Diana Navrátilová, Iñaki Odriozola, Sarah Piché-Choquette, Martina Štursová, Karel Švec, Vojtěch Tláskal, Michaela Urbanová, Lukáš Vlk, Jana Voříšková, Lucia Žifčáková, Petr Baldrian

## Abstract

Fungi are key players in vital ecosystem services, spanning carbon cycling, decomposition, symbiotic associations with cultural and wild plants and pathogenicity. The high importance of fungi in the ecosystem processes contrasts with the incompleteness of understanding of the patterns of fungal biogeography and the environmental factors that drive it. To close this gap of knowledge, we have here collected and validated data published on the composition of soil fungal communities in terrestrial environments including soil and plant-associated habitats and made them publicly accessible through a user interface at http://globalfungi.com. The GlobalFungi database contains over 650 million observations of fungal sequences across >20 000 samples with geographical locations and additional metadata contained in 207 original studies with millions of unique sequence variants of the fungal internal transcribed spacers (ITS) 1 and 2 representing fungal species and genera. As it is, the study represents the most comprehensive atlas of fungal distribution on the global scale open to further additions.

## Background & Summary

Fungi play fundamental roles in the ecosystem processes across all terrestrial biomes. As plant symbionts, pathogens or major decomposers of organic matter they substantially influence plant primary production, carbon mineralization and sequestration, and act as crucial regulators of the soil carbon balance^1,2^. The activities of fungi and their communities contribute to the production of clean water, food, and air and the suppression of disease-causing soil organisms. Soil biodiversity is thus increasingly recognized to provide services critical to food safety and human health^3^.

It is of high importance to determine how environmental factors affect the diversity and distribution of fungal communities. So far, only a few studies focused on fungal distribution and diversity on global scale^4–6^. Importantly, these single survey studies focused either on a limited number of biomes^4,5^, narrower taxonomical group within the Fungal kingdom^6^, or were restricted only to fungi inhabiting soil. While the benefit of a focused study is the fact that it can rely on standardized sample analysis and metadata collection, its limitation is the inability to cover large sampling efforts in space and time. On the other hand, since the advent of the high-throughput-sequencing methods, large amounts of sequencing data exist for fungi from terrestrial environments with sufficient metadata that allow their evaluation^7^. Recently, the meta-analysis of 36 papers reporting global diversity of soil fungi collected >3000 samples and helped to indicate that climate is an important factor for the global distribution of soil fungi and identify the hotspots of fungal diversity outside the tropics^8^. This approach clearly demonstrated the utility of a meta-study approach to address fungal biogeography, ecology and diversity. In addition, the compilation of these data demonstrated the fact that symbiotic mycorrhizal fungi that aid cultural and wild plants to access nutrients, are more likely to be affected by rapid changes of climate than other guilds of fungi, including plant pathogens^8^ and helped to identify which fungi tend to follow alien plants invading new environments^9^.

Here, we have undertaken a comprehensive collection and validation of data published on the composition of soil fungal communities in terrestrial environments including soil and plant-associated habitats. This approach enabled us to construct the GlobalFungi database containing, on March 16, 2020, over 120 million of unique sequence variants^10^ of the fungal internal transcribed spacers (ITS) 1 and 2, covering >20 000 samples contained in 207 original studies (Figure 1). The dataset of sequence variant frequencies across samples, accompanied by metadata retrieved from published papers and in global climate databases is made publicly available at http://globalfungi.com. To achieve the goal to make published data findable, accessible, interoperable and reusable, the user interface at the above address allows the users to search for individual sequences, fungal species hypotheses^11^, species or genera, to get a visual representation of their distribution in the environment and to access and download sequencing data and metadata. In addition, the user interface also allows authors to submit data from studies not yet covered and in this way to help to build the resource for the community of researchers in mycology, biogeography and ecology of fungi.

**Figure 1:**
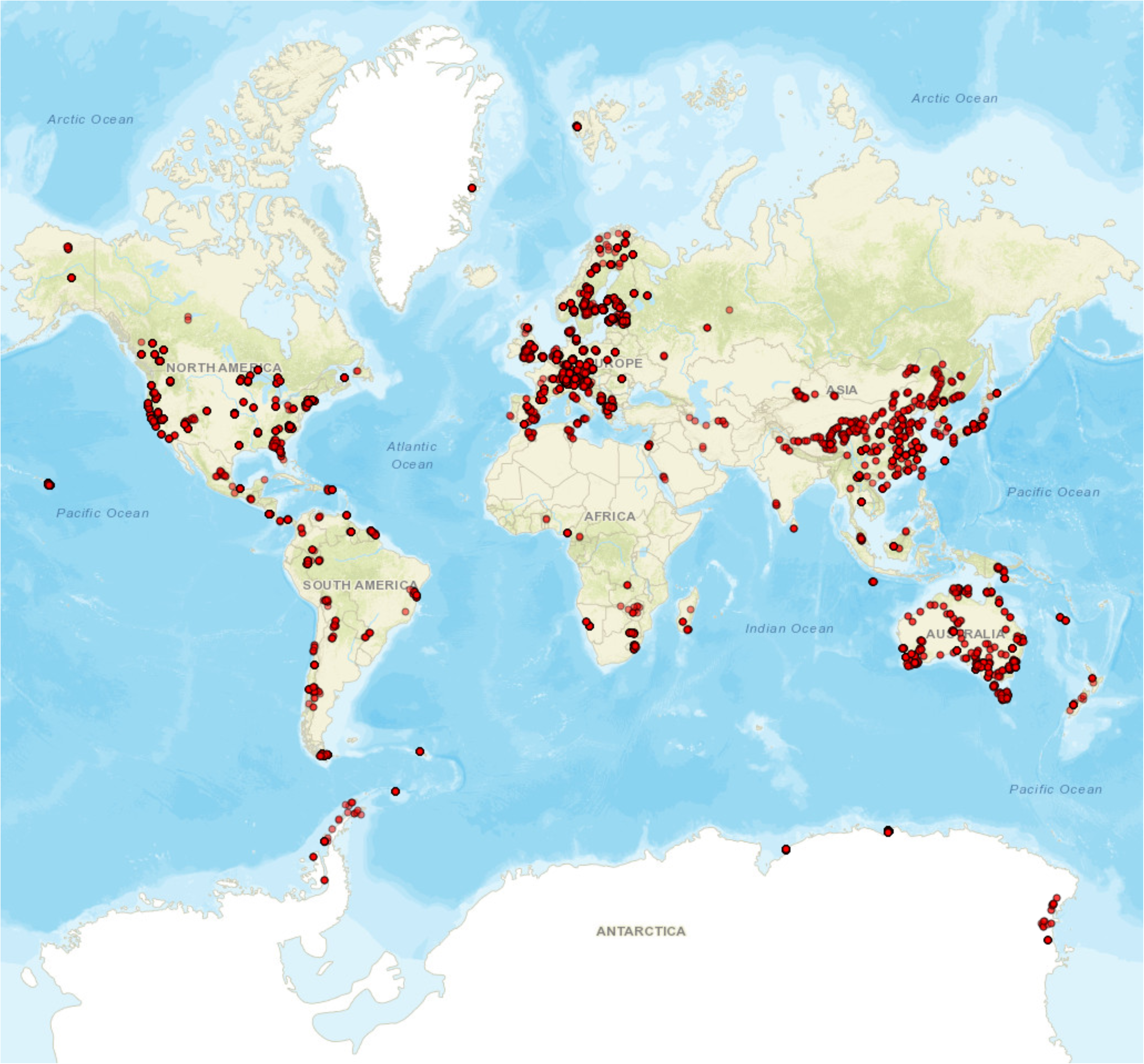
Map of locations of samples contained in the GlobalFungi database. Each point represents one or several samples where fungal community composition was reported using high-throughput-sequencing methods targeting the ITS1 or ITS2 marker of fungi. The map was created using the ‘leaflet’ package that uses an open-source JavaScript library for mobile-friendly interactive maps (GNU GENERAL PUBLIC LICENSE).

## Methods

### Data selection

We explored papers that used high-throughput sequencing for the analysis of fungal communities that were published until the beginning of 2019; in total, we explored 843 papers. The following selection criteria were used for the inclusion of samples (and, consequently, studies) into the dataset: (1) samples came from terrestrial biomes of soil, dead or live plant material (e.g., soil, litter, rhizosphere soil, topsoil, lichen, deadwood, root and shoot) and were not subject to experimental treatment; (2) the precise geographic location of each sample was recorded (GPS coordinates); (3) the whole fungal community was subject to amplicon sequencing (studies using group-specific primers were excluded); (4) the internal transcribed spacer regions (ITS1, ITS2, or both) were subject to amplification; (5) sequencing data (either in fasta or fastq format) were publicly available or provided by the authors of the study upon request, and the sequences were unambiguously assigned to samples; (6) the samples could be assigned to biomes according to the Environment Ontology (http://www.ontobee.org/ontology/ENVO)^8^. In total, there were 209 publications contained samples that matched our criteria (Online-Only Table 1).

#### Processing of sequencing data

For the processing of data, see Figure 2. Raw datasets from 207 studies, covering 20 009 individual samples were quality filtered by removing all sequences with the mean quality phred scores below 20. Each sequence was labelled using the combination of a sample ID and sequence ID, and the full ITS1 or ITS2 fungal region was extracted using Perl script ITSx v1.0.11^12^. ITS extraction resulted in a total of 416 291 533 full ITS1 and 231 278 756 full ITS2 sequences. The extracted ITS sequences were classified according to the representative sequence of the closest UNITE species hypothesis (SH) using BLASTN, considering the SH created using a 98.5% similarity threshold (general release 8.1 from 2.2.2019^11^. The sequence was classified to the best best hit SH only when the following thresholds were met: e-value <10e^−50^, sequence similarity >=98.5%. All representative sequences annotated as nonfungal were discarded. All representative sequences classified to any fungal SH and all unclassified sequences were used to build database sequence variant library. The number of sequence variants (unique nucleotide sequences) accessible through the database is 119 566 862.

**Figure 2:**
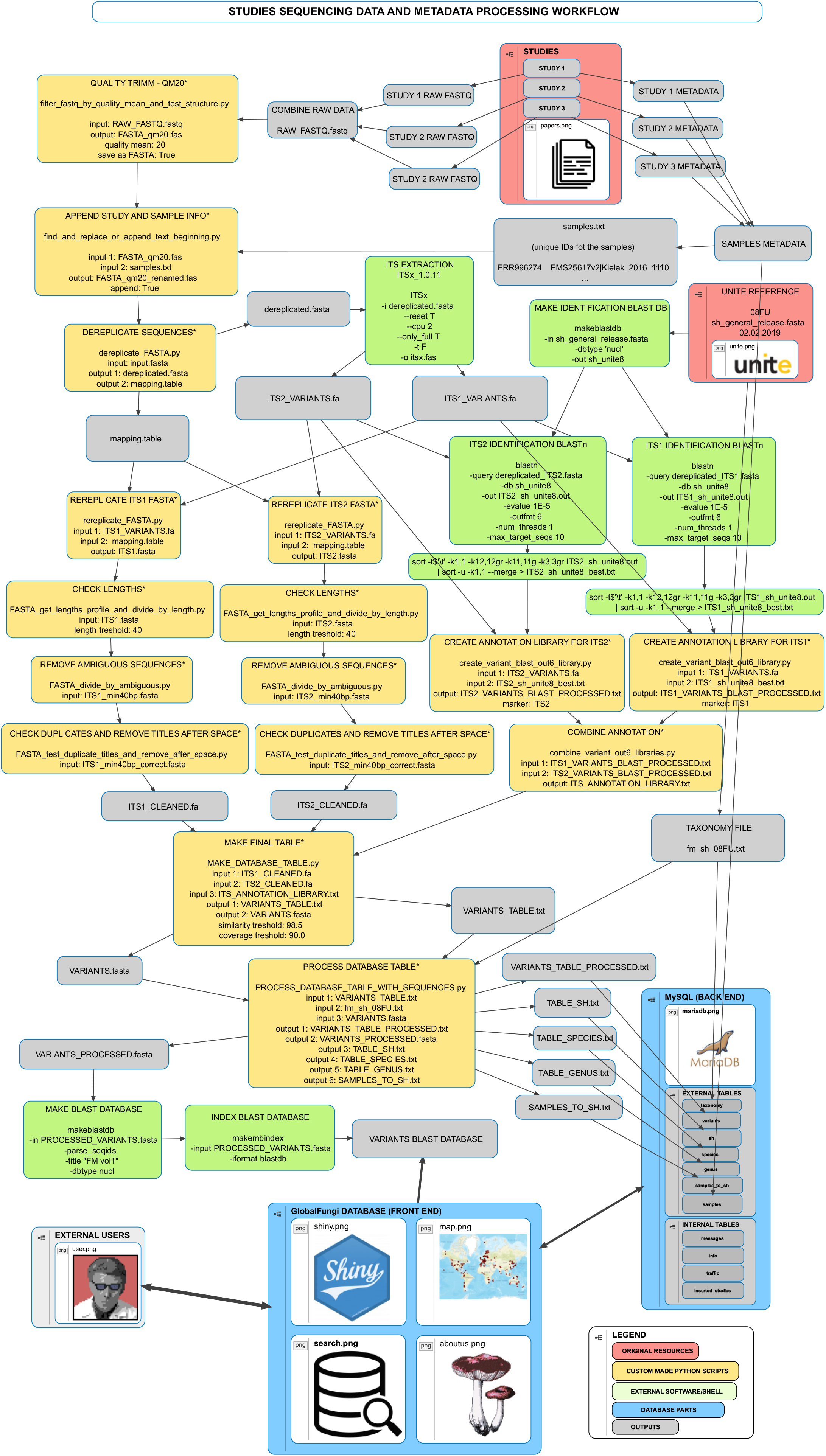
Processing of raw sequencing data for the GlobalFungi database. Workflow of processing of sequencing data included in the GlobalFungi database.

### Sample metadata

Sample metadata were collected from the published papers and/or public repositories where they were submitted by the authors. In some cases, metadata were obtained from the authors of individual studies upon request. The samples were assigned to continents, countries or specific locations, and all sites were categorized into biomes following the classification of Environment Ontology to a maximum achievable depth for each sample. The complete list of metadata included in the database is in Table 1.

**Table 1:** List of metadata contained in the GlobalFungi database. The table lists identifiers, units and sources of metadata contained in the database with the description of their content. Data source “original paper” may also represent additional metadata provided by the authors of the paper.

| Metadata identifier | Unit | Description of content | Source |
| --- | --- | --- | --- |
| Sample ID |  | unique identifier | generated |
| Longitude | degrees | Geographical longitude | original paper |
| Latitude | degrees | Geographical latitude | original paper |
| Continent |  | One of the following:<br>Africa/Antarctica/Asia/Australia/Europe/North America/South America | original paper |
| Sample type |  | One of the following: soil/rhizosphere<br>soil/litter/litter+humus/deadwood/lichen/shoot/root | original paper |
| Biome |  | One of the following: forest biome/woodland biome/shrubland biome/grassland biome/desert biome/tundra biome/mangrove biome/anthropogenic terrestrial biome/marine biome/freshwater biome/polar desert biome | original paper |
| Sampling year |  | Year of sample collection | original paper |
| Primers |  | Primers used | original paper |
| pH |  | pH | original paper |
| ITS total |  | Number of full ITS sequences extracted | generated |
| MAT (°C) | °C | Mean annual temperature from CHELSA database | CHELSA |
| MAP (mm) | mm | Mean annual precipitation from CHELSA database | CHELSA |

In addition to metadata provided by the authors of each study, we also extracted bioclimatic variables from the global CHELSA^13^ and WorldClim 2^14^ databases for each sample based on its GPS location. Since the results based on CHELSA and WorldClim 2 were comparable, we decided to include those from CHELSA, because precipitation patterns are better captured in the CHELSA dataset, in particular for mountain sites^13^.

For each sequence variant that was classified to SH, fungal species name and genus name was retrieved from the UNITE database^11^, when available.

## Data Records

The raw sequencing reads used to create the database are available at the following locations: One part of the raw ITS sequencing reads can be found from the National Center for Biotechnology Information (NCBI) Sequence Read Archive under accession numbers SRP001058^15^, SRP001175^16^, SRP006078^17^, SRP012868^18^, SRP013695^19^, SRP013944^20^, SRP015735^21^, SRP016090^22^, SRP026207^23^, SRP028404^24^, SRP033719^25^, SRP035356^26^, SRP040314^27^, SRP040786^28^, SRP041347^29^, SRP043106^30^, SRP043706^31^, SRP043982^32^, SRP044665^33^, SRP045166^34^, SRP045587^35^, SRP045746^36^, SRP045933^37^, SRP046049^38^, SRP048036^39^, SRP048856^40^, SRP049544^41^, SRP051033^42^, SRP052222^43^, SRP052716^44^, SRP055957^45^, SRP057433^46^, SRP057541^47^, SRP058508^48^, SRP058555^49^, SRP058851^50^, SRP059280^51^, SRP060838^52^, SRP061179^53^, SRP061305^54^, SRP061904^55^, SRP062647^56^, SRP063711^57^, SRP064158^58^, SRP065817^59^, SRP066030^60^, SRP066284^61^, SRP066331^62^, SRP067301^63^, SRP067367^64^, SRP068514^65^, SRP068608^66^, SRP068620^67^, SRP068654^68^, SRP069065^69^, SRP069742^70^, SRP070568^71^, SRP073070^72^, SRP073265^73^, SRP074055^74^, SRP074496^75^, SRP075989^76^, SRP079403^77^, SRP079521^78^, SRP080210^79^, SRP080428^80^, SRP080680^81^, SRP082472^82^, SRP082976^83^, SRP083394^84^, SRP083434^85^, SRP083901^86^, SRP087715^87^, SRP090261^88^, SRP090335^89^, SRP090490^90^, SRP090651^91^, SRP091741^92^, SRP091855^93^, SRP091867^94^, SRP092609^95^, SRP092777^96^, SRP093592^97^, SRP093928^98^, SRP094708^99^, SRP097883^100^, SRP101553^101^, SRP101605^102^, SRP102378^103^, SRP102417^104^, SRP102775^105^, SRP106137^106^, SRP106774^107^, SRP107174^108^, SRP107743^109^, SRP109164^110^, SRP109773^111^, SRP110522^112^, SRP110810^113^, SRP113348^114^, SRP114697^115^, SRP114821^116^, SRP115350^117^, SRP115464^118^, SRP115599^119^, SRP117302^120^, SRP118875^121^, SRP118960^122^, SRP119174^123^, SRP125864^124^, SRP132277^125^, SRP132591^126^, SRP132598^127^, SRP136886^128^, SRP139483^129^, SRP142723^130^, SRP148813^131^, SRP150527^132^, SRP151262^133^, SRP153934^134^, SRP160913^135^, SRP161632^136^ and SRP195764^137^, in the European Nucleotide Archive (ENA) Sequence Read Archive under accession numbers ERP001713^138^, ERP003251^139^, ERP003790^140^, ERP005177^141^, ERP005905^142^, ERP009341^143^, ERP010027^144^, ERP010084^145^, ERP010743^146^, ERP011924^147^, ERP012017^148^, ERP013208^149^, ERP013987^150^, ERP014227^151^, ERP017480^152^, ERP017851^153^, ERP017915^154^, ERP019580^155^, ERP019924^156^, ERP020657^157^, ERP022511^158^, ERP022742^159^, ERP023275^160^, ERP023718^161^, ERP023855^162^, ERP106131^163^, ERP107634^164^, ERP107636^165^, ERP110188^166^ and ERP112007^167^’, in the DNA Data Bank of Japan (DDBJ) under the accession numbers DRA000926^168^, DRA000937^169^, DRA001737^170^, DRA002424^171^, DRA002469^172^, DRA003024^173^, DRA003730^174^, DRA004913^175^, DRA006519^176^, DRP002783^177^, DRP003138^178^ and DRP005365^179^, in the Dryad Digital Repository under the DOIs https://doi.org/10.5061/dryad.2fc32^180^, http://dx.doi.org/10.5061/dryad.n82g9^181^, http://dx.doi.org/10.5061/dryad.2343k^182^, https://doi.org/10.5061/dryad.gp302^183^, http://dx.doi.org/10.5061/dryad.cq2rb^184^, https://doi.org/10.5061/dryad.8fn8j^185^ and https://doi.org/10.5061/dryad.216tp^186^, in the MG-RAST repository under the accession numbers 4783710.3^187^, 4702703.3^188^, 4524551.3^189^, 4544233.3^190^, 4683808.3^191^, 4684008.3^192^, 4696490.3^193^, 4715213.3^194^, 4563787.3^195^, 4563788.3^196^, 4620497.3^197^, 4620498.3^198^, mgp13293^199^, mgp1617^200^ and mgp9003^201^, in the PlutoF repository under the URLs https://plutof.ut.ee/#/filerepository/view/1561672^202^, https://plutof.ut.ee/#/filerepository/view/1562683^203^, https://plutof.ut.ee/#/doi/10.15156/BIO/100002^204^ and https://plutof.ut.ee/#/doi/10.15156/BIO/587446^205^, in the UNITE database under the URL http://unite.ut.ee/454_EcM_CMR.zip^206^, in GenBank under the accessions KAYV00000000.1^207^, KAYU00000000.1^208^, KAYT00000000.1^209^, SAMN02934078^210^ and SAMN02934079^211^, in the database of the Australian Antarctic Data Center under DOI http://dx.doi.org/10.4225/15/526f42ada05b1^212^, in the supplemental data of papers under URLs https://static-content.springer.com/esm/art%3A10.1038%2Fismej.2012.84/MediaObjects/41396_2012_BFismej201284_MOESM78_ESM.zip^213^ and https://static-content.springer.com/esm/art%3A10.1038%2Fismej.2015.238/MediaObjects/41396_2016_BFismej2015238_MOESM67_ESM.zip^214^, and on server of the Czech Academy of Sciences under URLs www.biomed.cas.cz/mbu/lbwrf/metastudy_datasets/GF0001.zip^215^, www.biomed.cas.cz/mbu/lbwrf/metastudy_datasets/GF0002.zip^216^, www.biomed.cas.cz/mbu/lbwrf/metastudy_datasets/GF0003.zip^217^, www.biomed.cas.cz/mbu/lbwrf/metastudy_datasets/GF0004.zip^218^, www.biomed.cas.cz/mbu/lbwrf/metastudy_datasets/GF0005.zip^219^, www.biomed.cas.cz/mbu/lbwrf/metastudy_datasets/GF0006.zip^220^, www.biomed.cas.cz/mbu/lbwrf/metastudy_datasets/GF0007.zip^221^, www.biomed.cas.cz/mbu/lbwrf/metastudy_datasets/GF0008.zip^222^, www.biomed.cas.cz/mbu/lbwrf/metastudy_datasets/GF0009.zip^223^, www.biomed.cas.cz/mbu/lbwrf/metastudy_datasets/GF0010.zip^224^, www.biomed.cas.cz/mbu/lbwrf/metastudy_datasets/GF0011.zip^225^, www.biomed.cas.cz/mbu/lbwrf/metastudy_datasets/GF0012.zip^226^.

The database contains two data types: sequence variants (individual nucleotide sequences) and samples. For each sequence variant, the following information is stored: sequence variant code, identification of samples where sequence variant occurs and the number of observations, the SH of best hit (when available), fungal species name (when available), fungal genus name (when available). For each sample, metadata information is stored (Table 1). Sequencing data and metadata are accessible at Figshare (GlobalFungi_ITS_variants.zip, GlobalFungi_metadata.xlsx). All database content is accessible using a public graphical user interface at http://globalfungi.com.

## Technical Validation

The technical validation included the screening of the data sources, sequencing data and data reliability. Regarding the data source screening, the data sources (published papers) were screened to fulfil the criteria outlined in the Methods section. The dataset was thoroughly checked for duplicities and for all records that appeared in multiple publications, only the first original publication of the dataset was considered as a datasource. Considering sequence quality, we have only utilized those primer pairs that are generally accepted to target general fungi^7,227^. Sequences were quality filtered by removing all sequences with the mean quality phred scores below 20 and sequences that did not represent complete ITS1 or ITS2 after extraction or those that were identified as chimeric by the ITS extraction software^12^ were removed. All representative sequences where the BLASTn search against the UNITE database^11^ resulted in a nonfungal organism, were discarded.

For data reliability, the geographic location represented by the GPS coordinates was validated first. For each sample set, the geographic location of the sample described in the text of the study was confronted with the location on the map. For those samples, where disagreement was recorded (e.g., terrestrial samples positioned in the ocean, location in another region than described in the text), the authors of each study were asked for correction. If impossible, such samples were deleted. The quality of sample metadata was checked and if they were outside the acceptable range (such as content of elements or organic matter >100%), these invalid metadata were removed.

## Usage Notes

The user interface at http://globalfungi.com enables the users to access the database in several ways (Figure 3). In taxon search, it is possible to seach for genera and species of fungi and for the 98.5% SH species hypotheses of UNITE, contained in the general release 8.1 from 2.2.2019. The search results open the options to download corresponding SH or corresponding sequence variants. It is also possible to view breakdown of samples by type, biome, mean annual temperature, mean annual precipitation, pH and continents. The results also contain an interactive map of the taxon distribution with relative abundances of sequences of the taxon across samples and a list of samples with metadata. Several modes of filtering of results are available as well.

**Figure 3:**
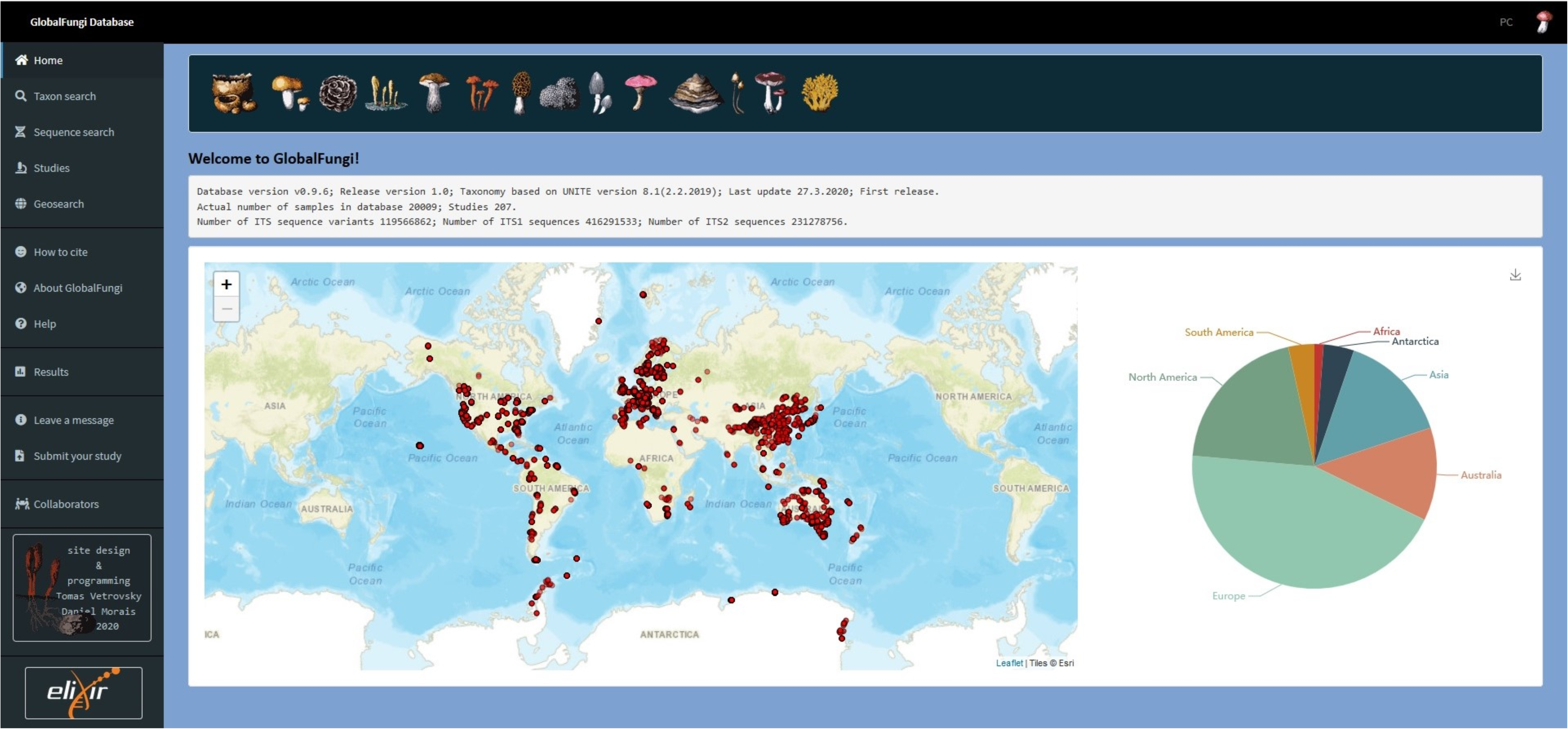
User interface to access the GlobalFungi database.

In sequence search, it is possible to search for multiple nucleotide sequences by exact match or BLAST retrieving the sequence variant best hit in the database, and alternatively, blasting a single sequence can deliver multiple ranked high score hits among the sequence variants. It is also possible to open individual studies and access their content and to select a group of samples on the map with a range of tools and retrieve data for these samples (such as the FASTA file with all occurring sequence variants).

Importantly, the database is intended to grow, both by the continuing activity of the authors and using the help of the scientific community. For that, web interface enabling the submission of studies not yet represented is available to users. The submission tool guides the submitting person through the steps where details about the publication, samples, sample metadata and sequences are sequentially collected. The submitted data will be added to the existing dataset after processing and validation by the authors.

## Code Availability

The workflow included several custom made python scripts (labelled by star) which are accessible here: https://github.com/VetrovskyTomas/GlobalFungi.

## Acknowledgements

We acknowledge funding from the Czech Science Foundation Grant No 18-26191S. ELIXIR CZ research infrastructure project LM2015047 by the Ministry of Education, Youth and Sports of the Czech Republic is acknowledged for hosting the database. All corresponding authors of published studies that provided additional information on the samples included in the database are gratefully acknowledged.

## Author contributions

T.V., P.K. and P.B. jointly conceived the study. C.L. and coordinated data acquisition, T.V., D.M., M.K. and P.B. designed the database, T.V. and D.M. developed the database and created the user interface. C.A.G., S.A.H., B.D.B., K.B., V.B., F.D., R.Z.H., M.J., J.K., C.L., S.L., R.L.M., T.Mar., T.Maš., L.Me., L.Mi., T.Mi., S.M., D.N., I.O., S.P.C., M.Š., K.Š., V.T., M.U., L.V., J.V. and L.Ž. identified data sources, processed and analysed sequencing information, collated and analysed metadata. P.B. drafted the manuscript along with T.V., C.L. and P.K. All authors wrote and reviewed the manuscript.

T.V., D.M., P.K. and C.L. contributed equally to this work.

---

**Online-only Table 1:**
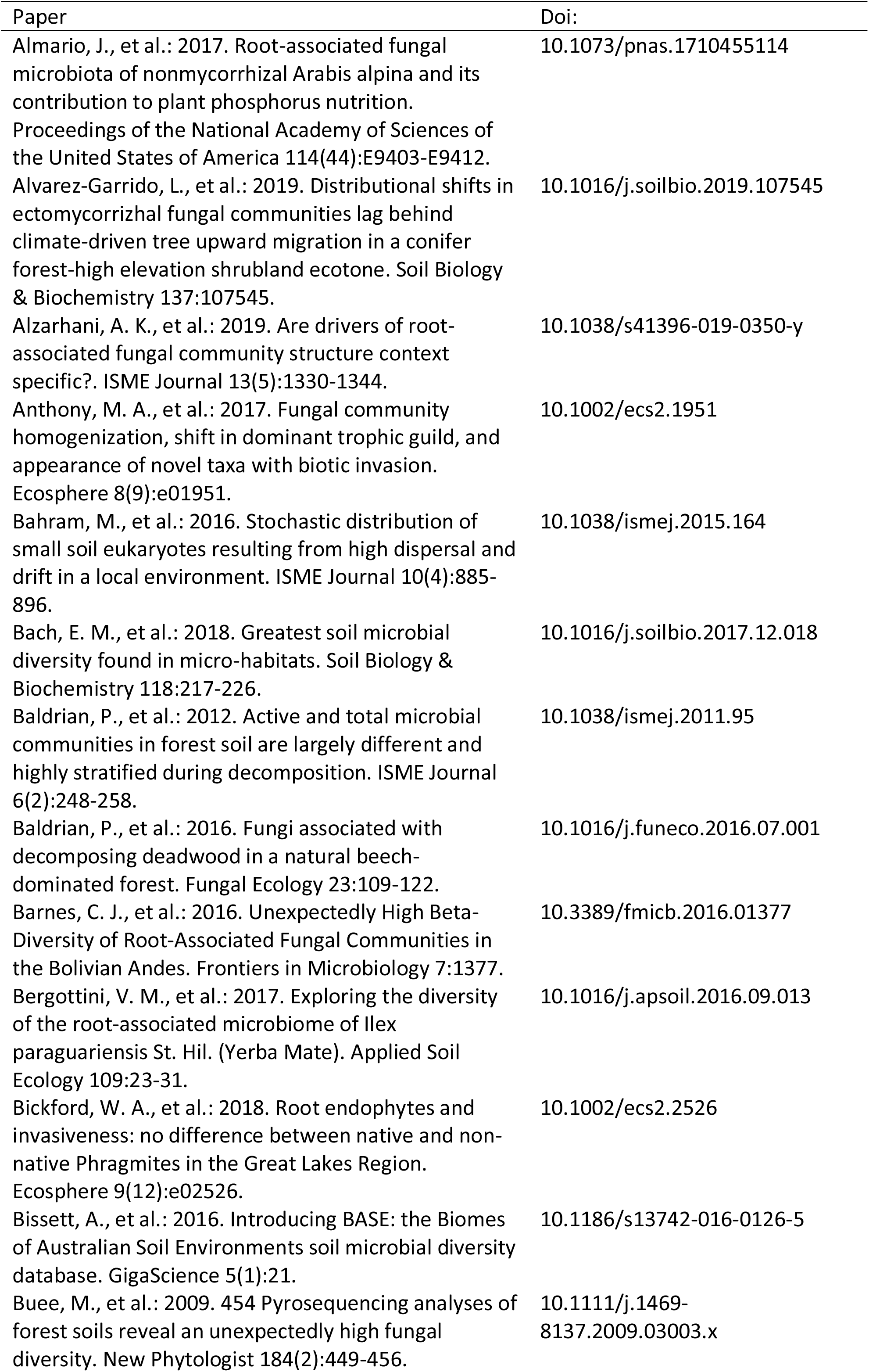
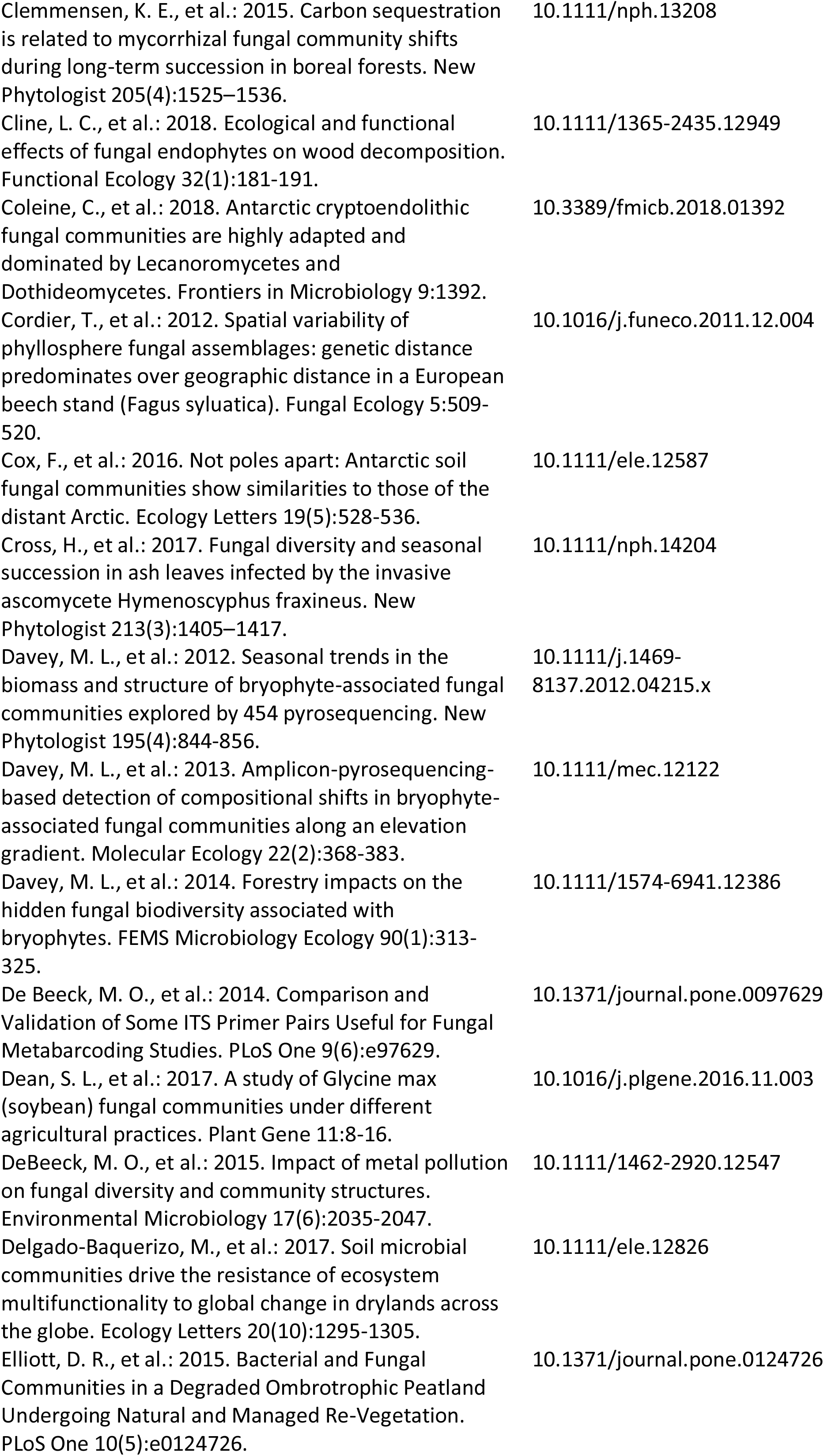
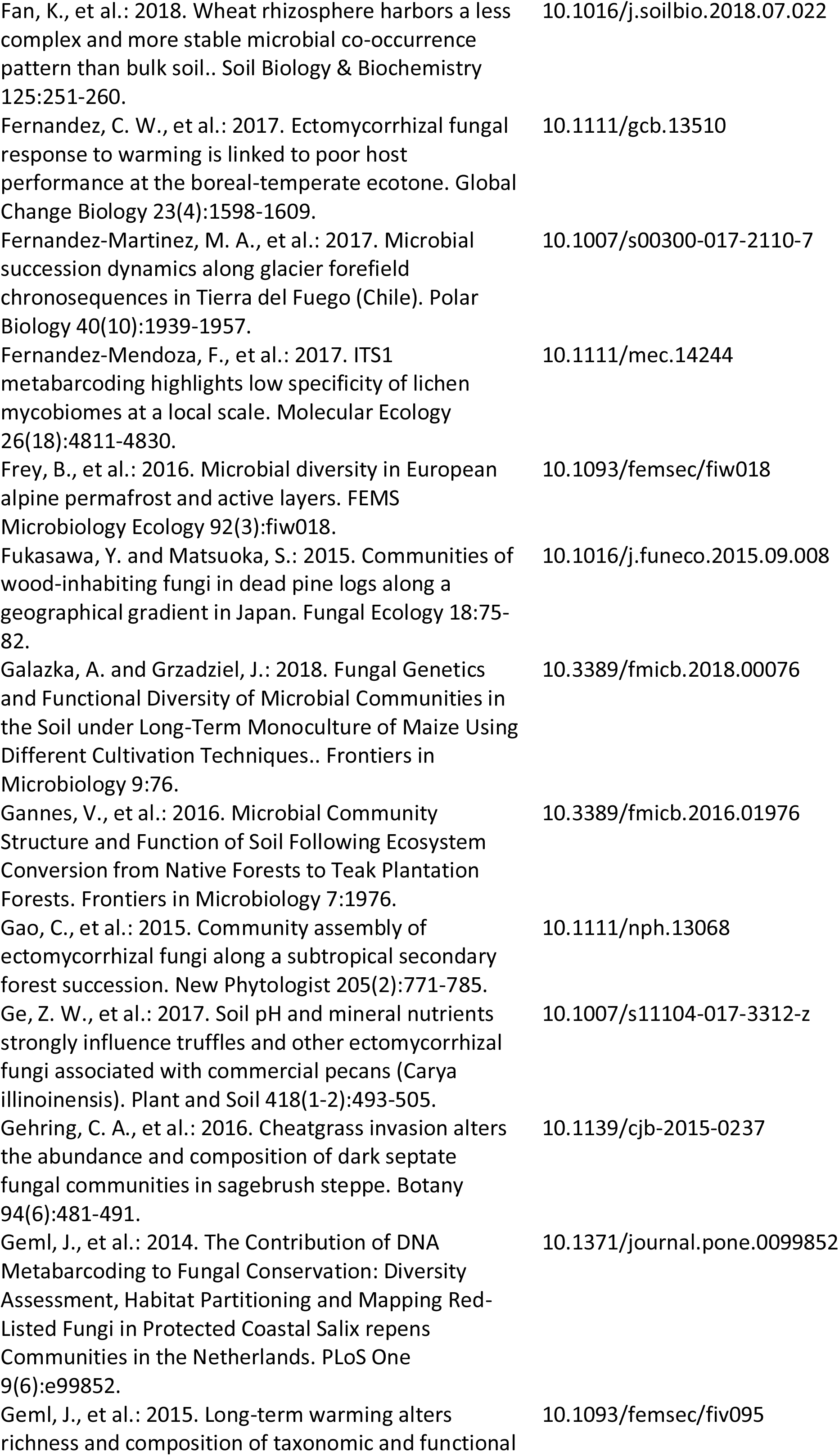
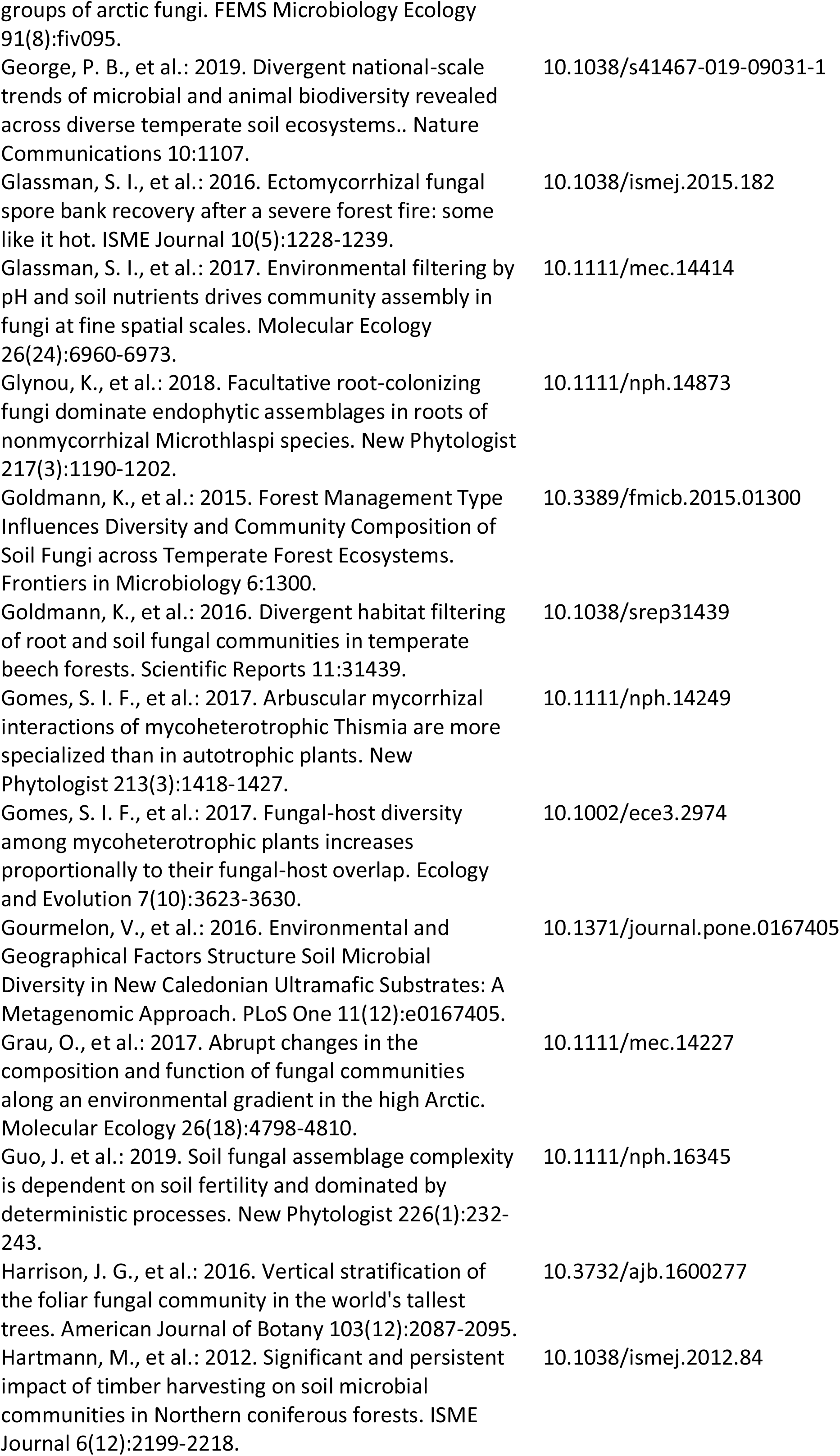
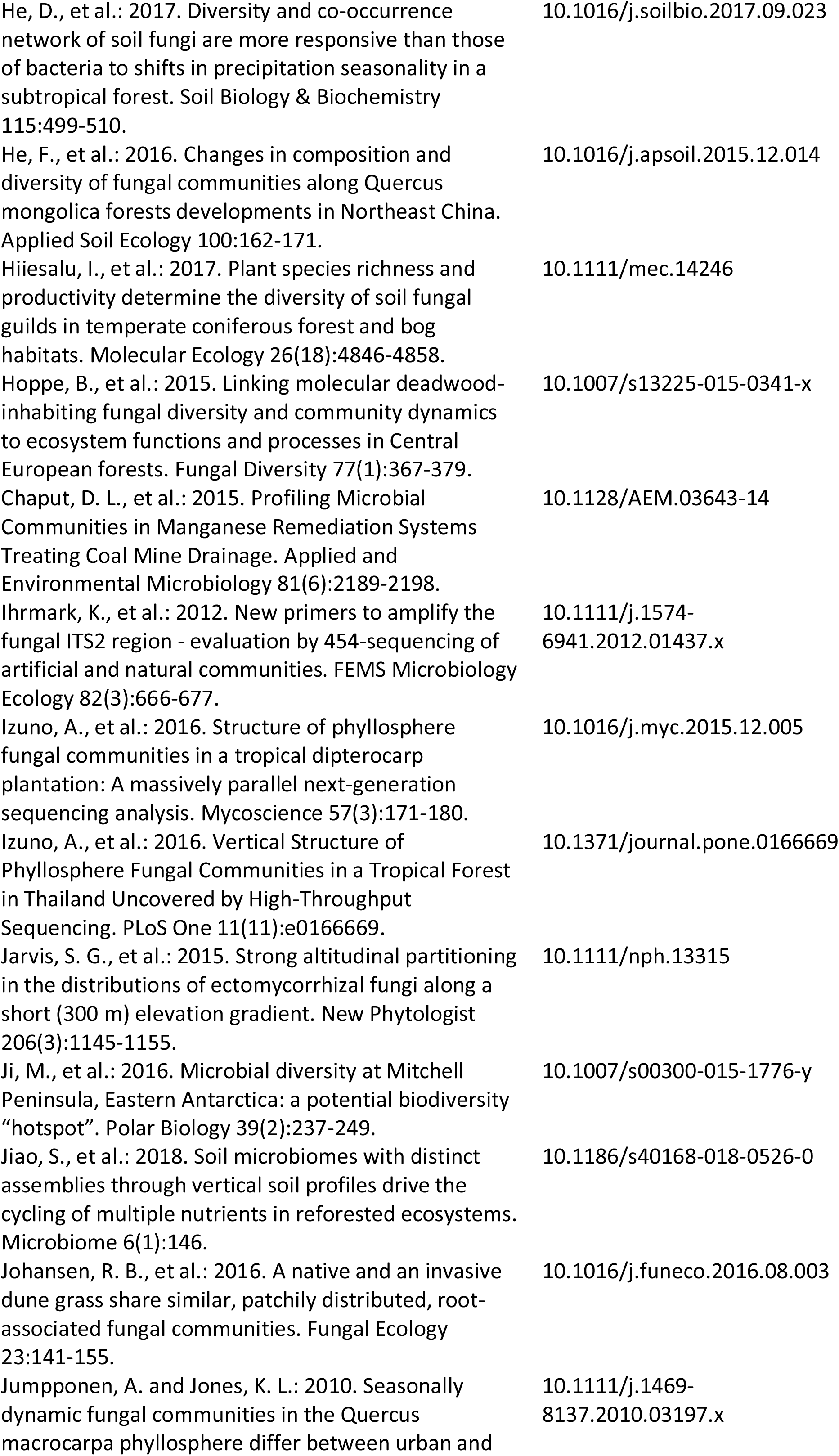
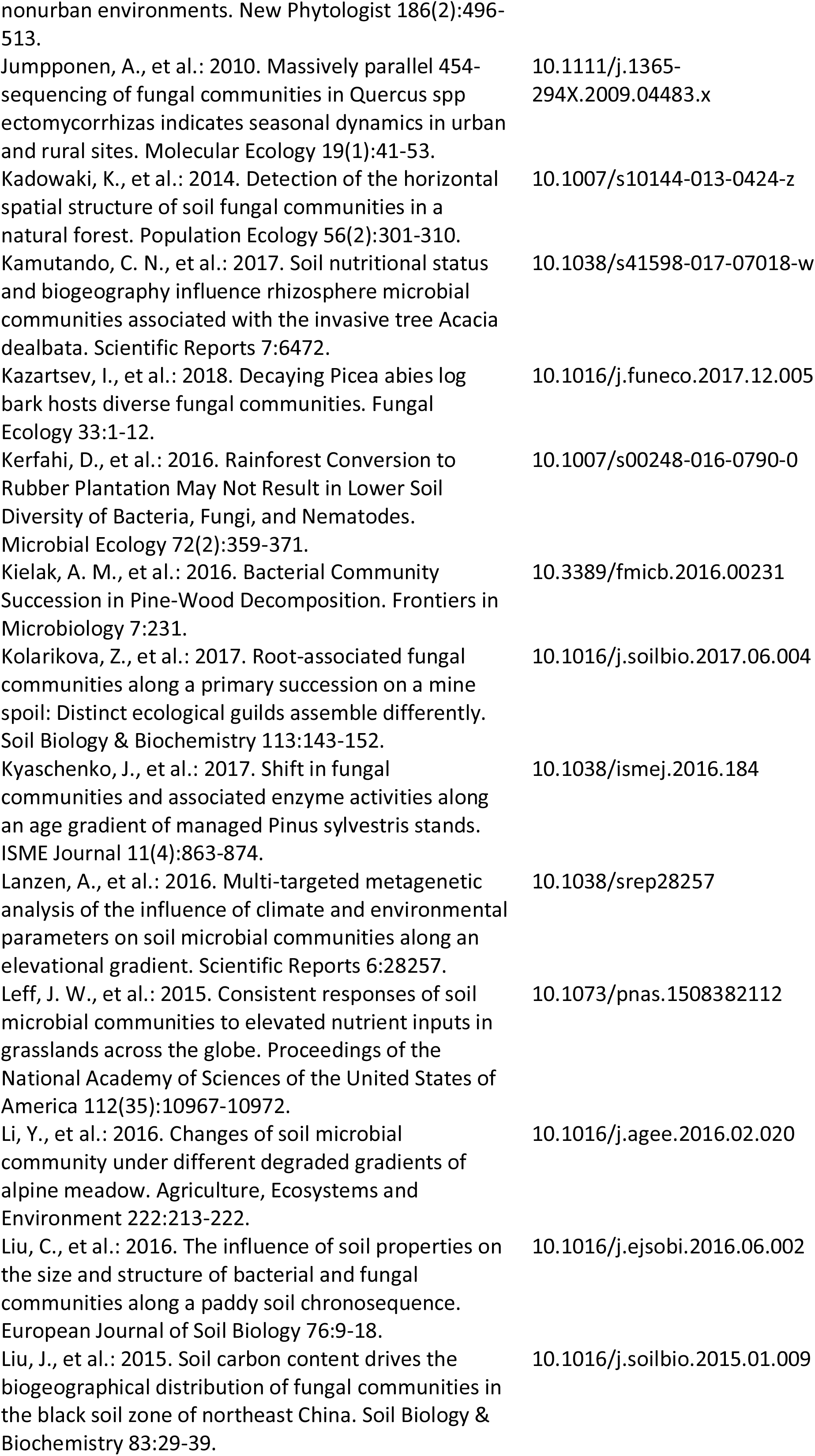
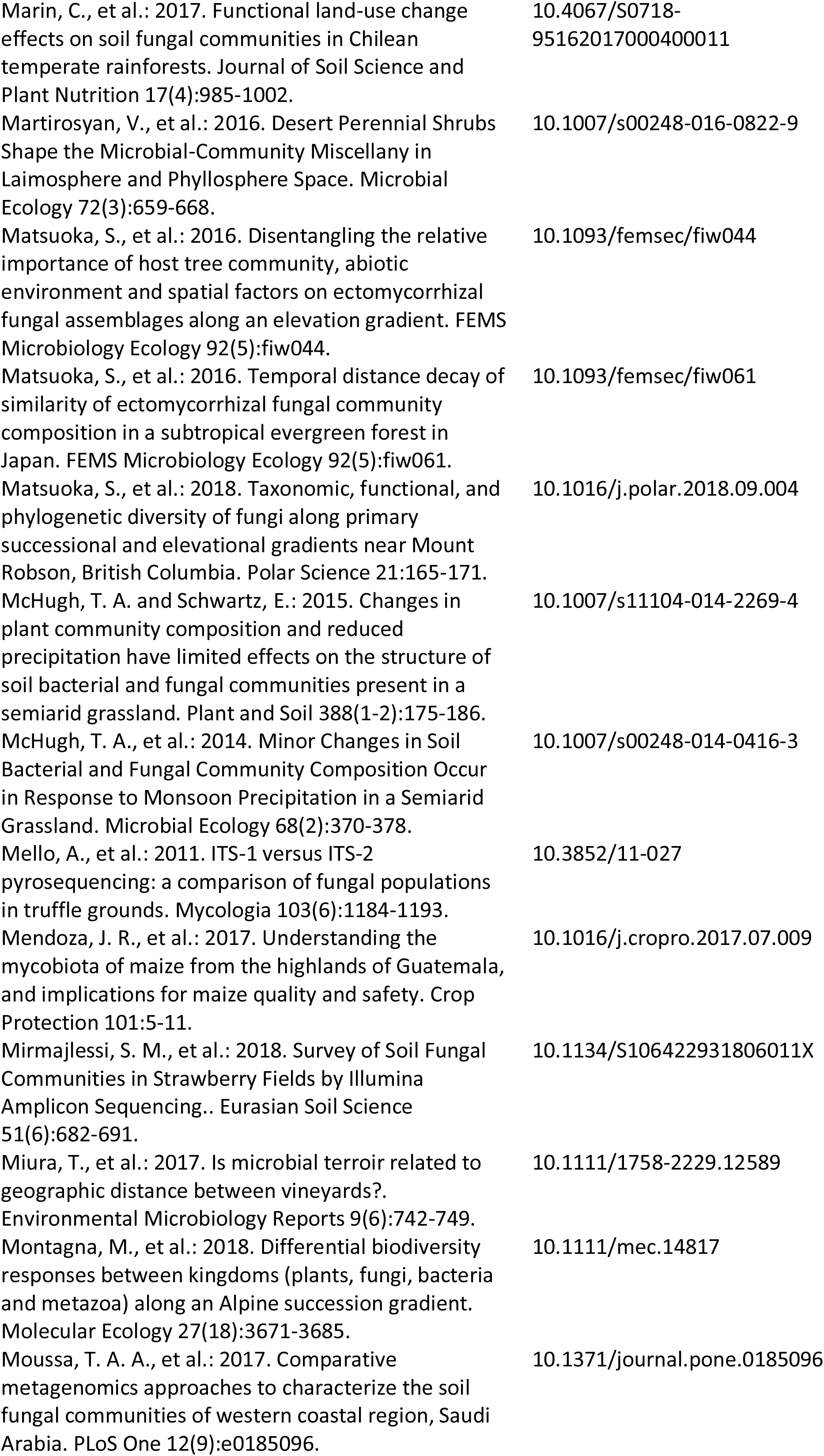
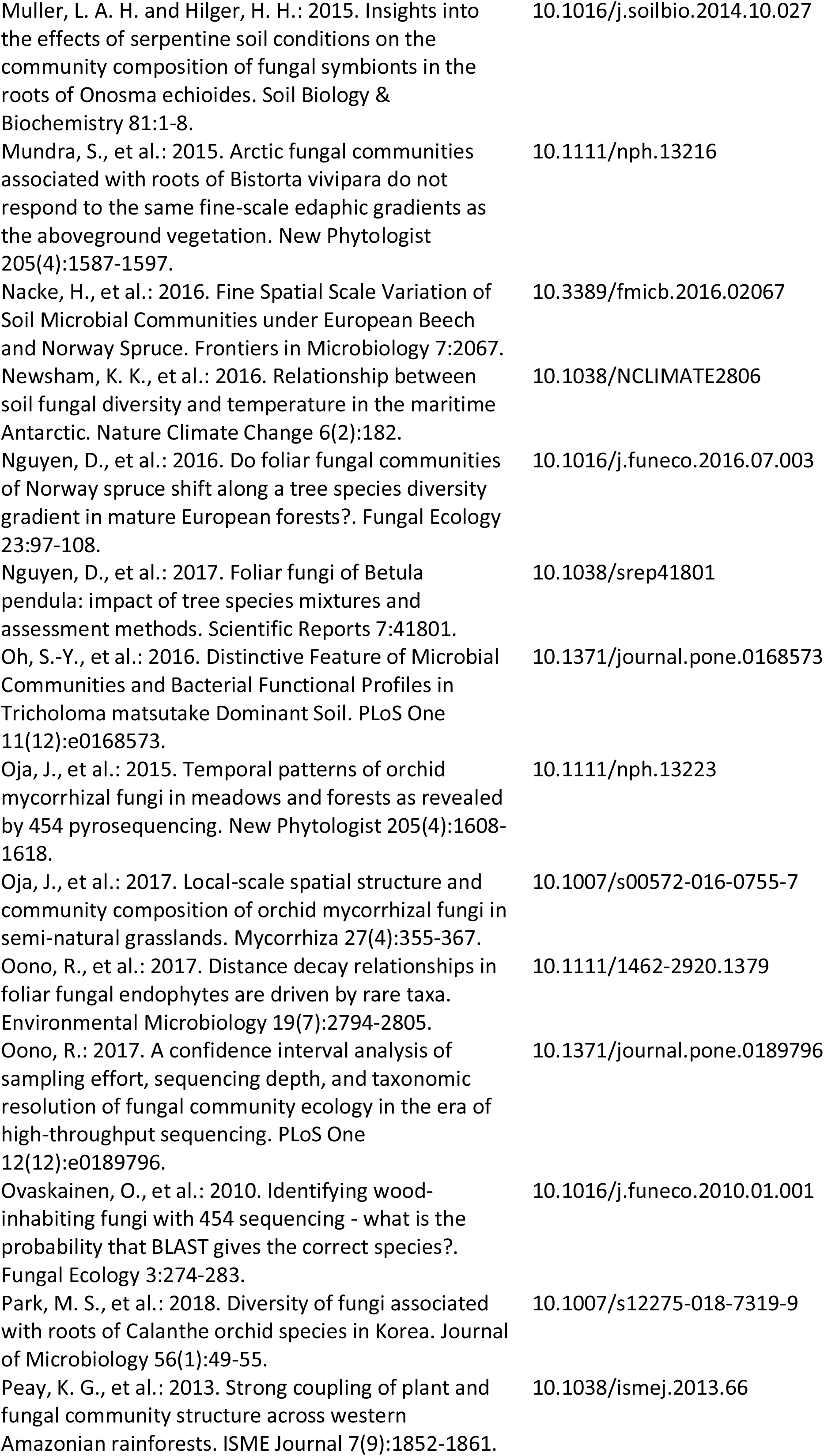
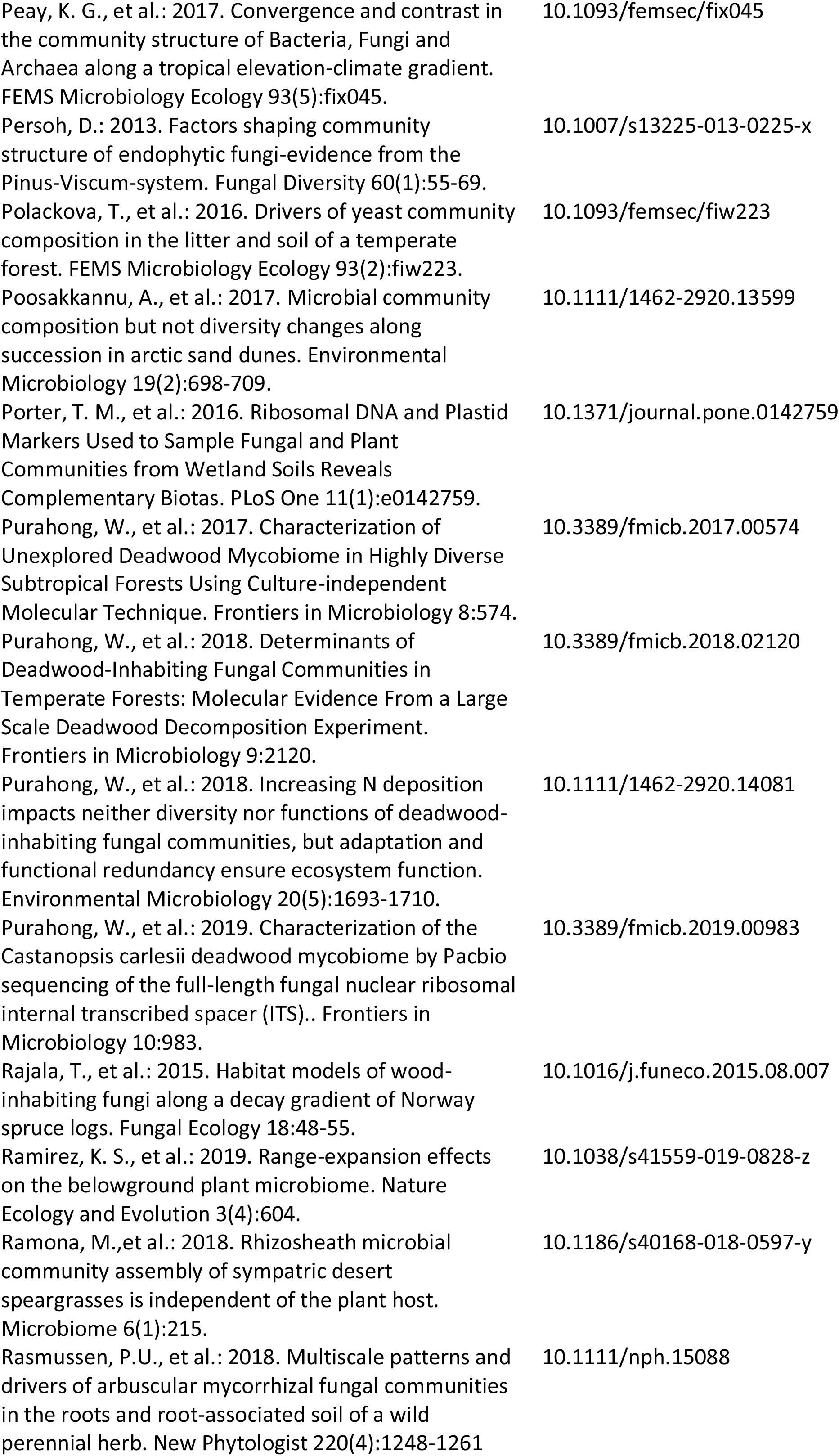
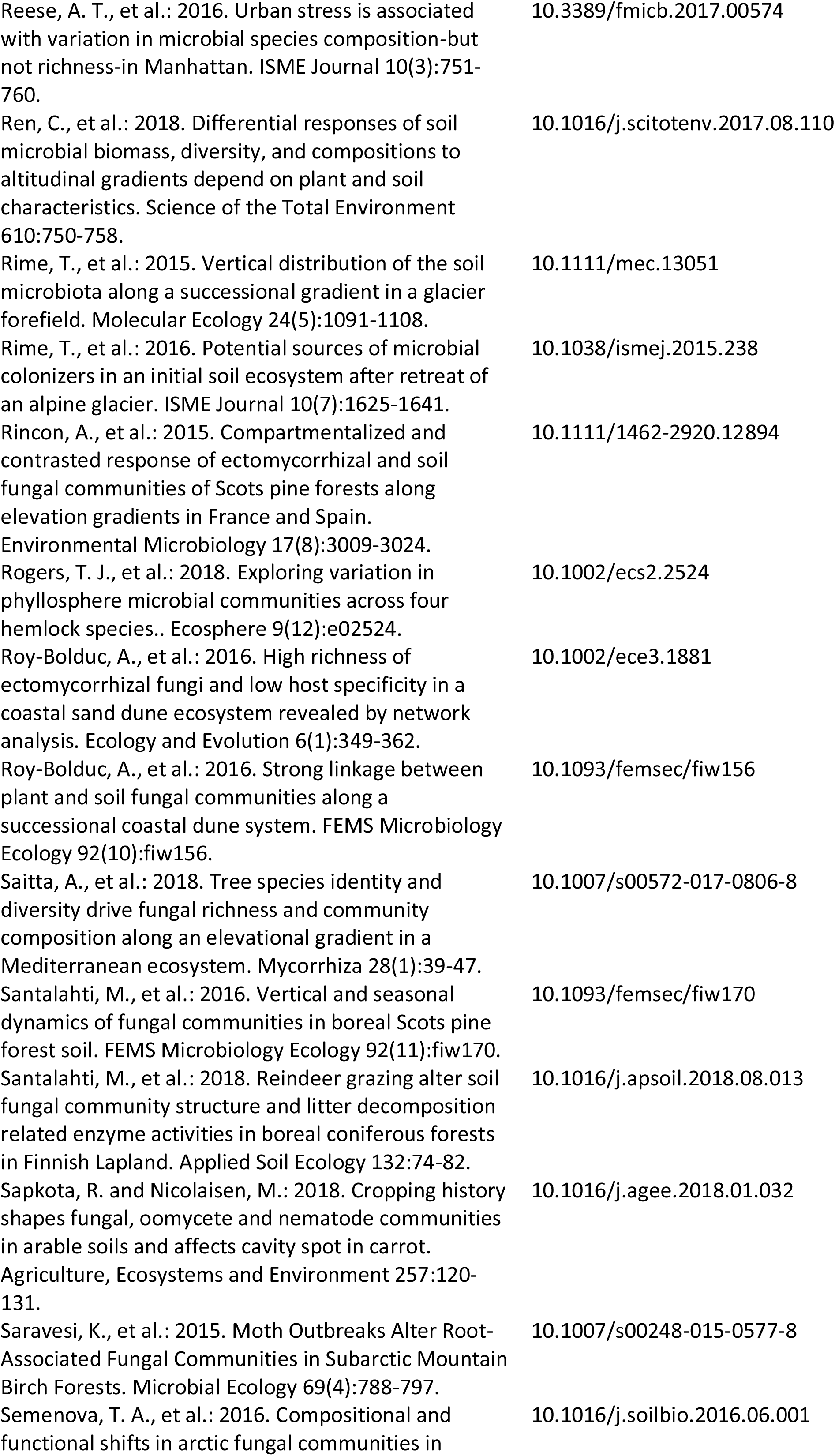
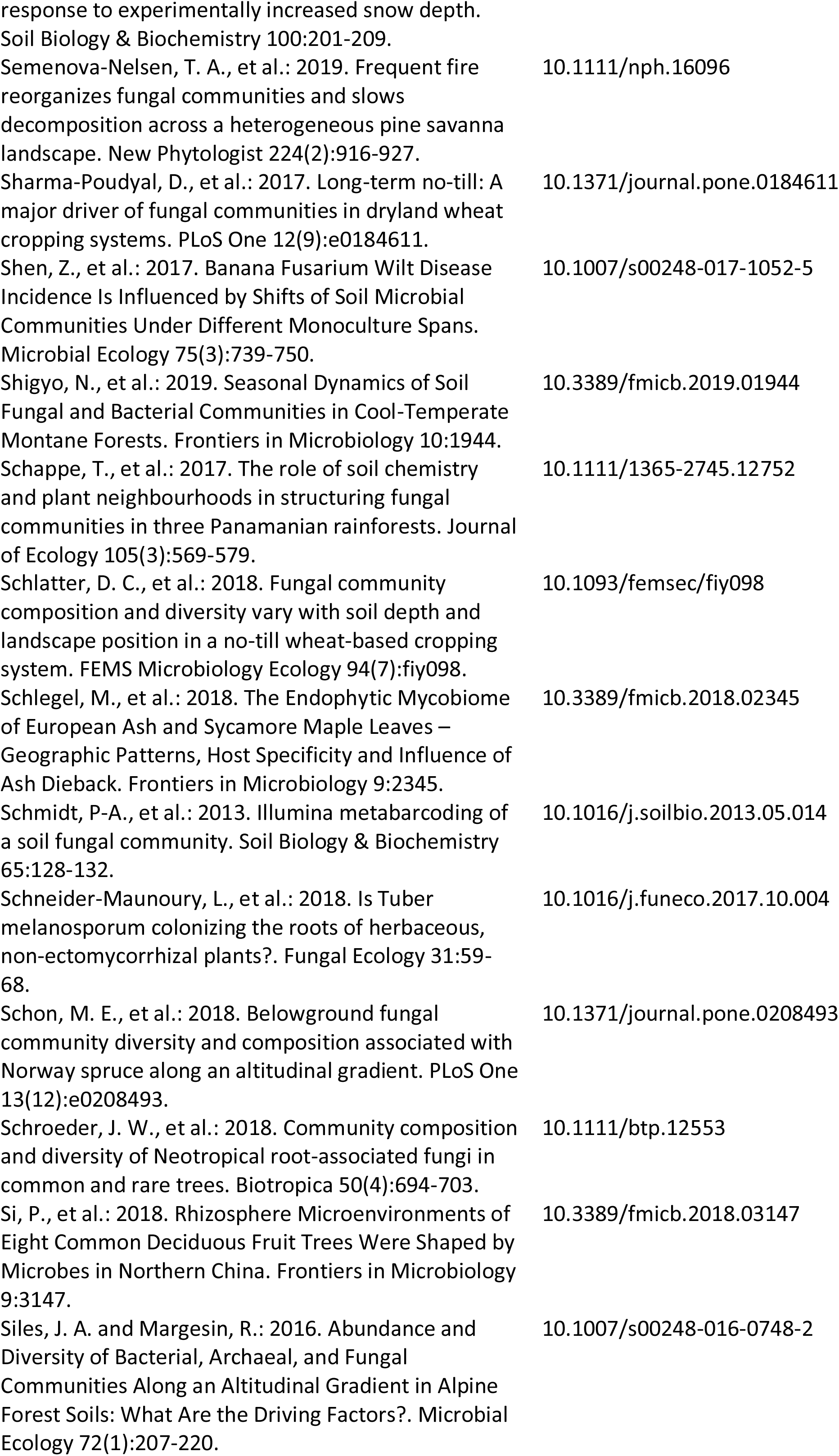
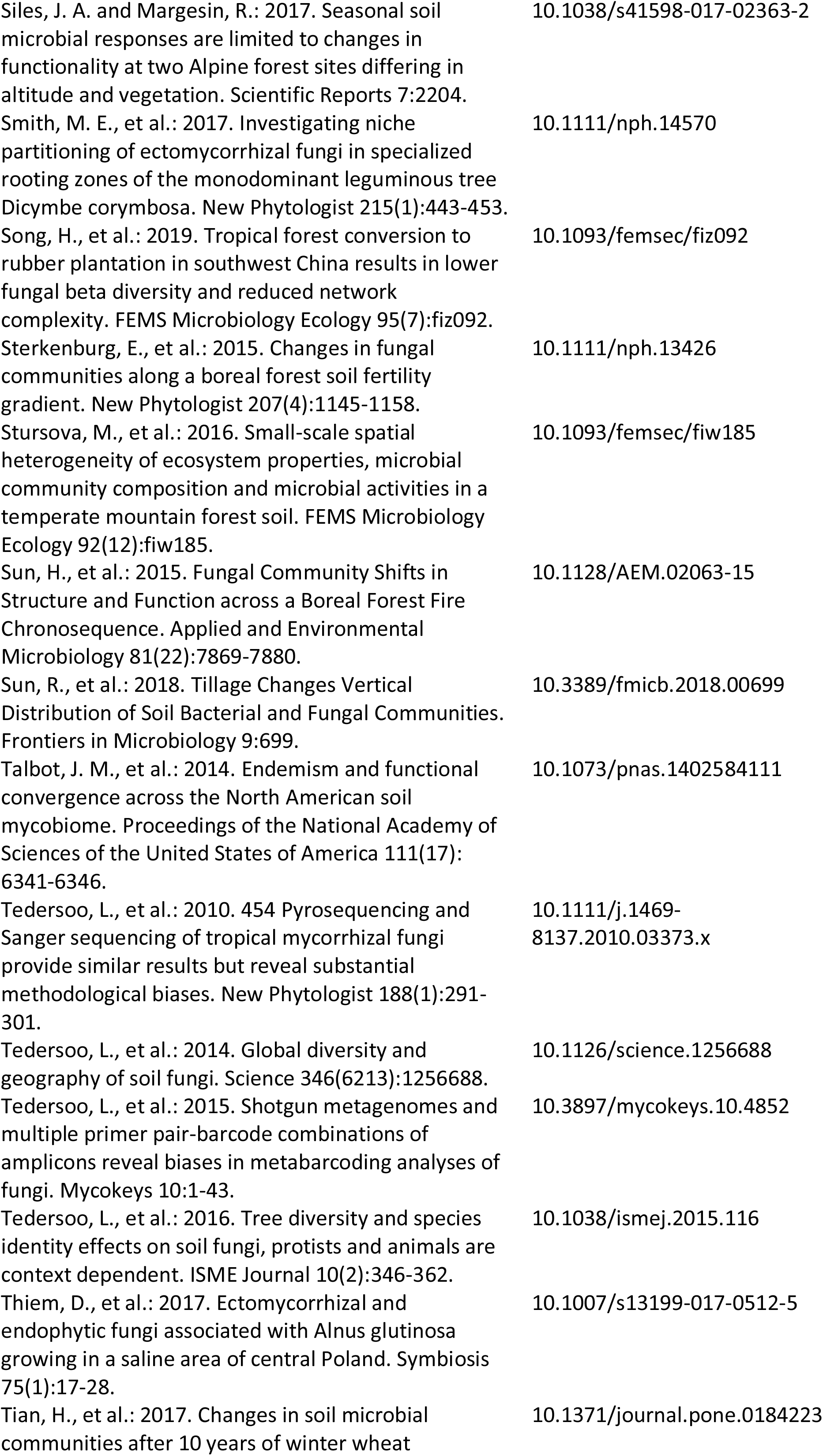
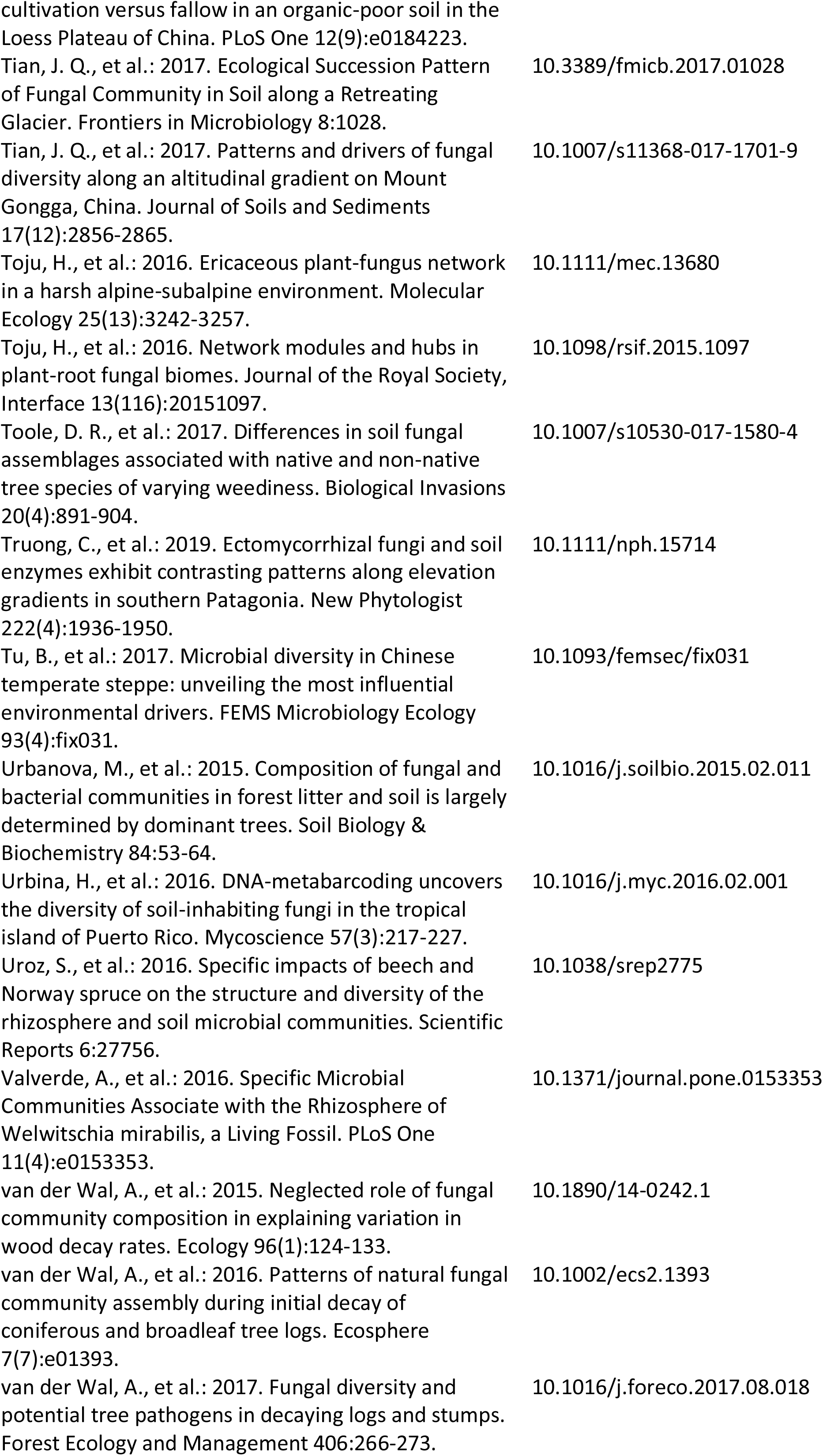
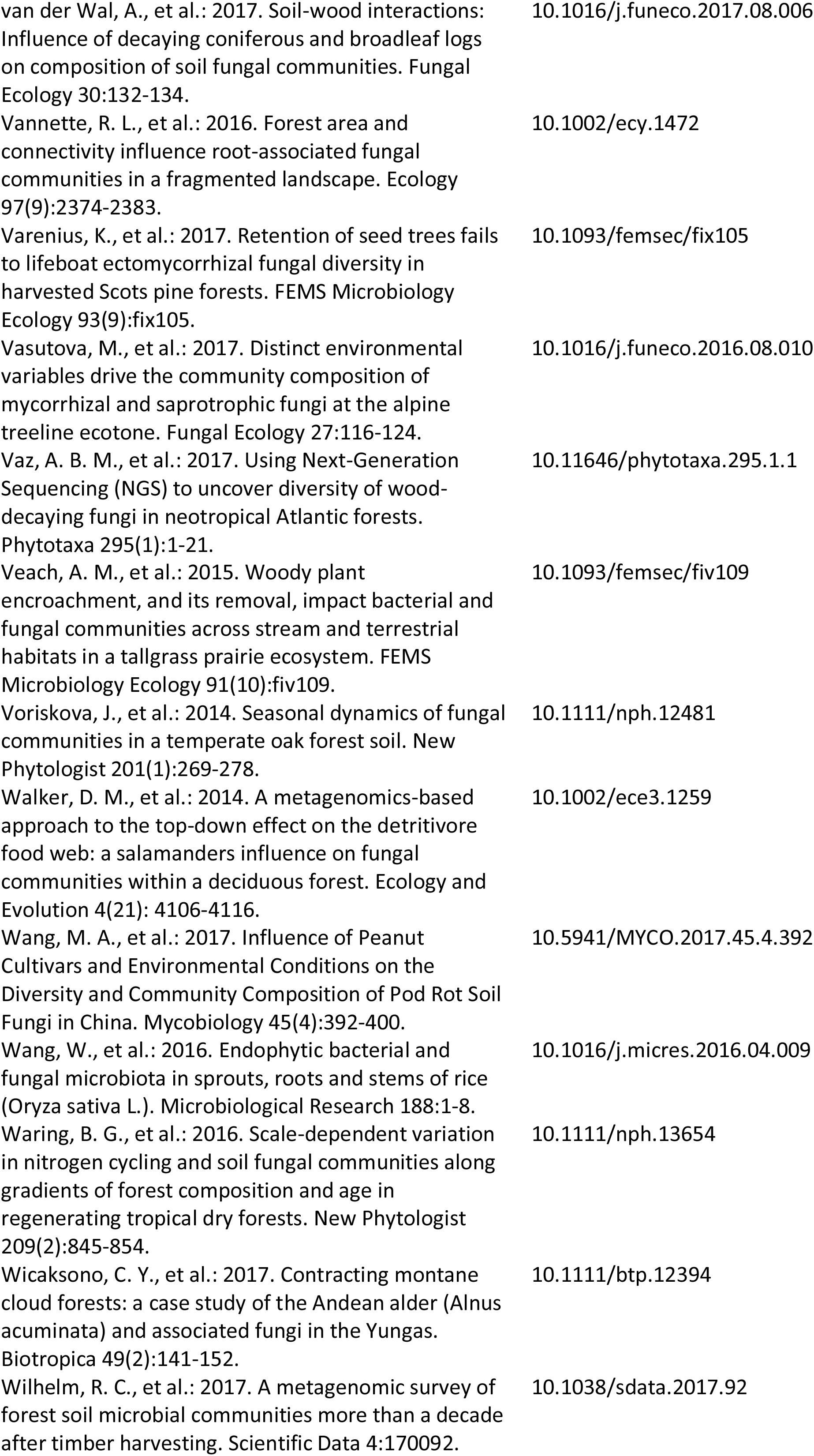
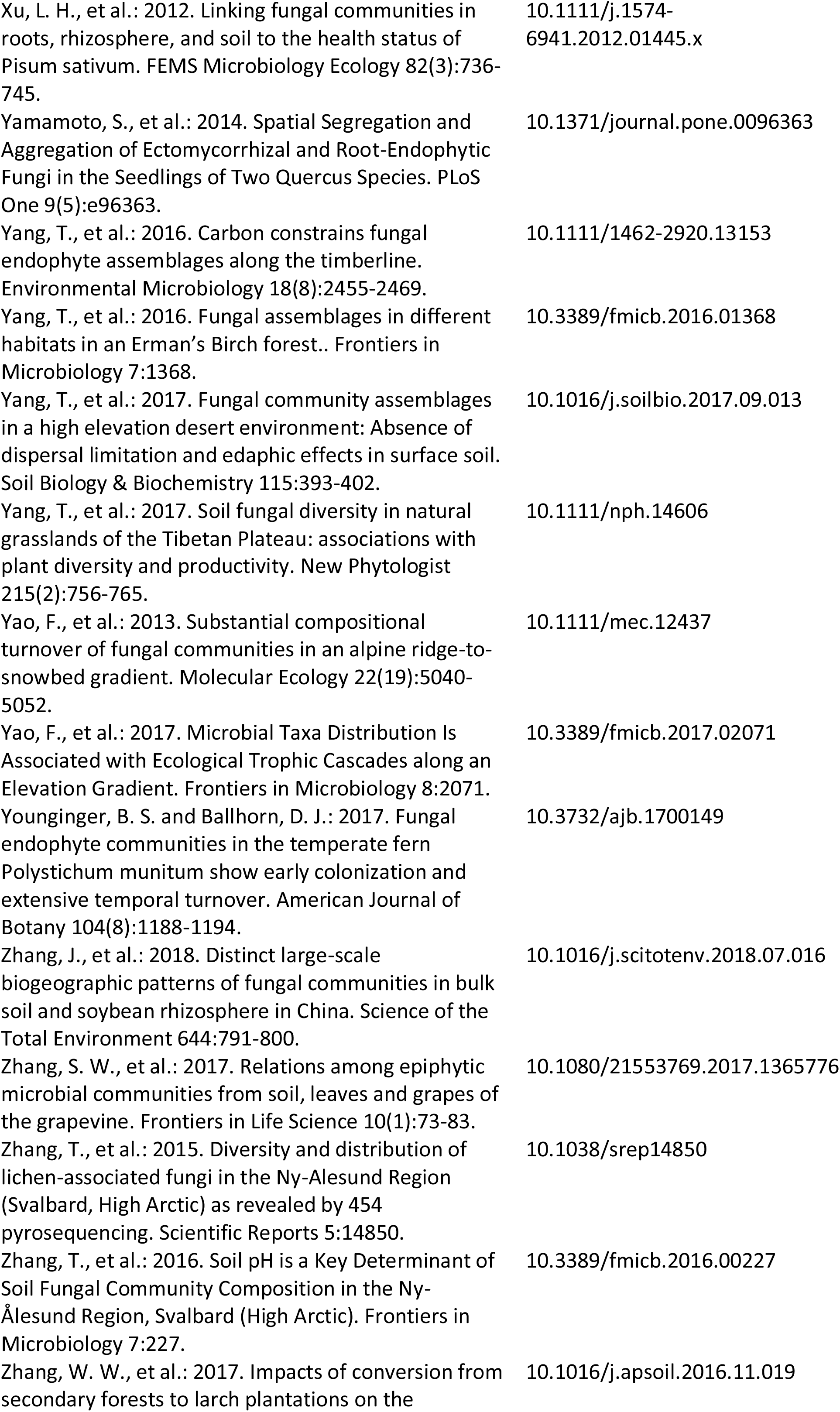
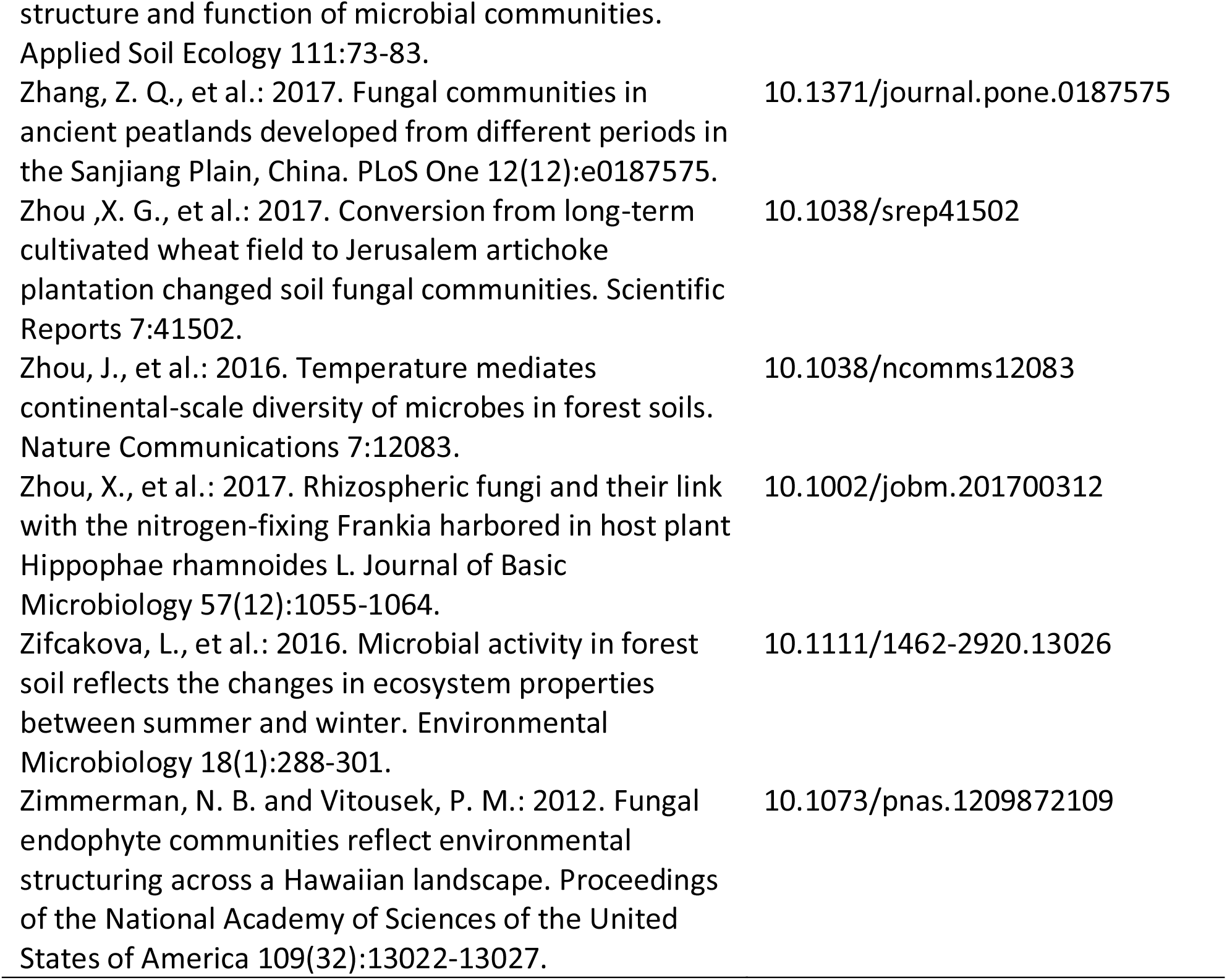
List of papers that were used as data sources for GlobalFungi database with digital object identifiers.

## References

1. Crowther, T.W., Todd-Brown, K.E.O., Rowe, C.W., Wieder, W.R., Carey, J.C., Machmuller, M.B., Snoek, B.L., Fang, S., Zhou, G., Allison, S.D., Blair, J.M., Bridgham, S.D., Burton, A.J., Carrillo, Y., Reich, P.B., Clark, J.S., Classen, A.T., Dijkstra, F.A., Elberling, B., Emmett, B.A., Estiarte, M., Frey, S.D., Guo, J., Harte, J., Jiang, L., Johnson, B.R., Kroel-Dulay, G., Larsen, K.S., Laudon, H., Lavallee, J.M., Luo, Y., Lupascu, M., Ma, L.N., Marhan, S., Michelsen, A., Mohan, J., Niu, S., Pendall, E., Penuelas, J., Pfeifer-Meister, L., Poll, C., Reinsch, S., Reynolds, L.L., Schmidt, I.K., Sistla, S., Sokol, N.W., Templer, P.H., Treseder, K.K., Welker, J.M. & Bradford, M.A. Quantifying global soil carbon losses in response to warming. Nature 540, 104–108 (2016).

2. Peay, K.G., Kennedy, P.G. & Talbot, J.M. Dimensions of biodiversity in the Earth mycobiome. Nature Rev. Microbiol. 14, 434–447 (2016).

3. Wall, D.H., Nielsen, U.N. & Six, J. Soil biodiversity and human health. Nature 528, 69–76 (2015).

4. Tedersoo, L., Bahram, M., Polme, S., Koljalg, U., Yorou, N.S., Wijesundera, R., Ruiz, L.V., Vasco-Palacios, A.M., Thu, P.Q., Suija, A., Smith, M.E., Sharp, C., Saluveer, E., Saitta, A., Rosas, M., Riit, T., Ratkowsky, D., Pritsch, K., Poldmaa, K., Piepenbring, M., Phosri, C., Peterson, M., Parts, K., Partel, K., Otsing, E., Nouhra, E., Njouonkou, A.L., Nilsson, R.H., Morgado, L.N., Mayor, J., May, T.W., Majuakim, L., Lodge, D.J., Lee, S.S., Larsson, K.H., Kohout, P., Hosaka, K., Hiiesalu, I., Henkel, T.W., Harend, H., Guo, L.D., Greslebin, A., Grelet, G., Geml, J., Gates, G., Dunstan, W., Dunk, C., Drenkhan, R., Dearnaley, J., De Kesel, A., Dang, T., Chen, X., Buegger, F., Brearley, F.Q., Bonito, G., Anslan, S., Abell, S. & Abarenkov, K. Global diversity and geography of soil fungi. Science 346, 1256688 (2014).

5. Bahram, M., Hildebrand, F., Forslund, S.K., Anderson, J.L., Soudzilovskaia, N.A., Bodegom, P.M., Bengtsson-Palme, J., Anslan, S., Coelho, L.P., Harend, H., Huerta-Cepas, J., Medema, M.H., Maltz, M.R., Mundra, S., Olsson, P.A., Pent, M., Polme, S., Sunagawa, S., Ryberg, M., Tedersoo, L. & Bork, P. Structure and function of the global topsoil microbiome. Nature 560, 233–237 (2018).

6. Egidi, E., Delgado-Baquerizo, M., Plett, J.M., Wang, J., Eldridge, D.J., Bardgett, R.D., Maestre, F.T. & Singh, B.K. A few Ascomycota taxa dominate soil fungal communities worldwide. Nature Commun. 10, 2369 (2019).

7. Nilsson, R.H., Anslan, S., Bahram, M., Wurzbacher, C., Baldrian, P. & Tedersoo, L. Mycobiome diversity: high-throughput sequencing and identification of fungi. Nature Rev. Microbiol. 17, 95–109 (2019).

8. Větrovský, T., Kohout, P., Kopecký, M., Machac, A., Man, M., Bahnmann, B.D., Brabcová, V., Choi, J., Meszárošová, L., Human, Z.R., Lepinay, C., Lladó, S., Lopez-Mondejar, R., Martinovic, T., Mašínová, T., Morais, D., Navrátilová, D., Odriozola, I., Štursová, M., Švec, K., Tláskal, V., Urbanová, M., Wan, J., Žifčáková, L., Howe, A., Ladau, J., Peay, K.G., Storch, D., Wild, J. & Baldrian, P. A meta-analysis of global fungal distribution reveals climate-driven patterns. Nature Commun. 10, 5142 (2019).

9. Vlk, L., Tedersoo, L., Antl, T., Větrovský, T., Abarenkov, K., Pergl, J., Albrechtová, J., Vosátka, M., Baldrian, P., Pyšek, P. & Kohout, P. Early successional ectomycorrhizal fungi are more likely to naturalize outside their native range than other ectomycorrhizal fungi. New Phytol. in press, doi: 10.1111/nph.16557.

10. Thompson, L.R., Sanders, J.G., McDonald, D., Amir, A., Ladau, J., Locey, K.J., Prill, R.J., Tripathi, A., Gibbons, S.M., Ackermann, G., Navas-Molina, J.A., Janssen, S., Kopylova, E., Vázquez-Baeza, Y., González, A., Morton, J.T., Mirarab, S., Zech Xu, Z., Jiang, L., Haroon, M.F., Kanbar, J., Zhu, Q., Jin Song, S., Kosciolek, T., Bokulich, N.A., Lefler, J., Brislawn, C.J., Humphrey, G., Owens, S.M., Hampton-Marcell, J., Berg-Lyons, D., McKenzie, V., Fierer, N., Fuhrman, J.A., Clauset, A., Stevens, R.L., Shade, A., Pollard, K.S., Goodwin, K.D., Jansson, J.K., Gilbert, J.A. Knight, R. & The Earth Microbiome Project, C. A communal catalogue reveals Earth’s multiscale microbial diversity. Nature 551, 457–463 (2017).

11. Nilsson, R.H., Larsson, K.H., Taylor, A.F.S., Bengtsson-Palme, J., Jeppesen, T.S., Schigel, D., Kennedy, P., Picard, K., Glockner, F.O., Tedersoo, L., Saar, I., Koljalg, U. & Abarenkov, K. The UNITE database for molecular identification of fungi: handling dark taxa and parallel taxonomic classifications. Nucleic Acids Res. 47, D259–D264 (2019).

12. Bengtsson-Palme, J., Ryberg, M., Hartmann, M., Branco, S., Wang, Z., Godhe, A., De Wit, P., Sanchez-Garcia, M., Ebersberger, I., de Sousa, F., Amend, A.S., Jumpponen, A., Unterseher, M., Kristiansson, E., Abarenkov, K., Bertrand, Y.J.K., Sanli, K., Eriksson, K.M., Vik, U., Veldre, V. & Nilsson, R.H. Improved software detection and extraction of ITS1 and ITS2 from ribosomal ITS sequences of fungi and other eukaryotes for analysis of environmental sequencing data. Meth. Ecol. Evol. 4, 914–919 (2013).

13. Karger, D.N., Conrad, O., Bohner, J., Kawohl, T., Kreft, H., Soria-Auza, R.W., Zimmermann, N.E., Linder, H.P. & Kessler, M. Data Descriptor: Climatologies at high resolution for the earth's land surface areas. Scientific Data 4, 170122 (2017).

14. Fick, S.E. & Hijmans, R.J. WorldClim 2: new 1-km spatial resolution climate surfaces for global land areas. International J. Climatol. 37, 4302–4315 (2017).

15. NCBI Sequence Read Archive, https://identifiers.org/ncbi/insdc.sra:SRP001058 (2010).

16. NCBI Sequence Read Archive, https://identifiers.org/ncbi/insdc.sra:SRP001175 (2010).

17. NCBI Sequence Read Archive, https://identifiers.org/ncbi/insdc.sra:SRP006078 (2011).

18. NCBI Sequence Read Archive, https://identifiers.org/ncbi/insdc.sra:SRP012868 (2012).

19. NCBI Sequence Read Archive, https://identifiers.org/ncbi/insdc.sra:SRP013695 (2012).

20. NCBI Sequence Read Archive, https://identifiers.org/ncbi/insdc.sra:SRP013944 (2016).

21. NCBI Sequence Read Archive, https://identifiers.org/ncbi/insdc.sra:SRP015735 (2015).

22. NCBI Sequence Read Archive, https://identifiers.org/ncbi/insdc.sra:SRP016090 (2015).

23. NCBI Sequence Read Archive, https://identifiers.org/ncbi/insdc.sra:SRP026207 (2014).

24. NCBI Sequence Read Archive, https://identifiers.org/ncbi/insdc.sra:SRP028404 (2015).

25. NCBI Sequence Read Archive, https://identifiers.org/ncbi/insdc.sra:SRP033719 (2015).

26. NCBI Sequence Read Archive, https://identifiers.org/ncbi/insdc.sra:SRP035356 (2015).

27. NCBI Sequence Read Archive, https://identifiers.org/ncbi/insdc.sra:SRP040314 (2014).

28. NCBI Sequence Read Archive, https://identifiers.org/ncbi/insdc.sra:SRP040786 (2015).

29. NCBI Sequence Read Archive, https://identifiers.org/ncbi/insdc.sra:SRP041347 (2015).

30. NCBI Sequence Read Archive, https://identifiers.org/ncbi/insdc.sra:SRP043106 (2015).

31. NCBI Sequence Read Archive, https://identifiers.org/ncbi/insdc.sra:SRP043706 (2017).

32. NCBI Sequence Read Archive, https://identifiers.org/ncbi/insdc.sra:SRP043982 (2015).

33. NCBI Sequence Read Archive, https://identifiers.org/ncbi/insdc.sra:SRP044665 (2016).

34. NCBI Sequence Read Archive, https://identifiers.org/ncbi/insdc.sra:SRP045166 (2015).

35. NCBI Sequence Read Archive, https://identifiers.org/ncbi/insdc.sra:SRP045587 (2016).

36. NCBI Sequence Read Archive, https://identifiers.org/ncbi/insdc.sra:SRP045746 (2014).

37. NCBI Sequence Read Archive, https://identifiers.org/ncbi/insdc.sra:SRP045933 (2015).

38. NCBI Sequence Read Archive, https://identifiers.org/ncbi/insdc.sra:SRP046049 (2016).

39. NCBI Sequence Read Archive, https://identifiers.org/ncbi/insdc.sra:SRP048036 (2016).

40. NCBI Sequence Read Archive, https://identifiers.org/ncbi/insdc.sra:SRP048856 (2015).

41. NCBI Sequence Read Archive, https://identifiers.org/ncbi/insdc.sra:SRP049544 (2015).

42. NCBI Sequence Read Archive, https://identifiers.org/ncbi/insdc.sra:SRP051033 (2016).

43. NCBI Sequence Read Archive, https://identifiers.org/ncbi/insdc.sra:SRP052222 (2017).

44. NCBI Sequence Read Archive, https://identifiers.org/ncbi/insdc.sra:SRP052716 (2015).

45. NCBI Sequence Read Archive, https://identifiers.org/ncbi/insdc.sra:SRP055957 (2015).

46. NCBI Sequence Read Archive, https://identifiers.org/ncbi/insdc.sra:SRP057433 (2016).

47. NCBI Sequence Read Archive, https://identifiers.org/ncbi/insdc.sra:SRP057541 (2016).

48. NCBI Sequence Read Archive, https://identifiers.org/ncbi/insdc.sra:SRP058508 (2016).

49. NCBI Sequence Read Archive, https://identifiers.org/ncbi/insdc.sra:SRP058555 (2016).

50. NCBI Sequence Read Archive, https://identifiers.org/ncbi/insdc.sra:SRP058851 (2018).

51. NCBI Sequence Read Archive, https://identifiers.org/ncbi/insdc.sra:SRP059280 (2016).

52. NCBI Sequence Read Archive, https://identifiers.org/ncbi/insdc.sra:SRP060838 (2016).

53. NCBI Sequence Read Archive, https://identifiers.org/ncbi/insdc.sra:SRP061179 (2016).

54. NCBI Sequence Read Archive, https://identifiers.org/ncbi/insdc.sra:SRP061305 (2017).

55. NCBI Sequence Read Archive, https://identifiers.org/ncbi/insdc.sra:SRP061904 (2015).

56. NCBI Sequence Read Archive, https://identifiers.org/ncbi/insdc.sra:SRP062647 (2016).

57. NCBI Sequence Read Archive, https://identifiers.org/ncbi/insdc.sra:SRP063711 (2017).

58. NCBI Sequence Read Archive, https://identifiers.org/ncbi/insdc.sra:SRP064158 (2017).

59. NCBI Sequence Read Archive, https://identifiers.org/ncbi/insdc.sra:SRP065817 (2017).

60. NCBI Sequence Read Archive, https://identifiers.org/ncbi/insdc.sra:SRP066030 (2016).

61. NCBI Sequence Read Archive, https://identifiers.org/ncbi/insdc.sra:SRP066284 (2017).

62. NCBI Sequence Read Archive, https://identifiers.org/ncbi/insdc.sra:SRP066331 (2017).

63. NCBI Sequence Read Archive, https://identifiers.org/ncbi/insdc.sra:SRP067301 (2017).

64. NCBI Sequence Read Archive, https://identifiers.org/ncbi/insdc.sra:SRP067367 (2016).

65. NCBI Sequence Read Archive, https://identifiers.org/ncbi/insdc.sra:SRP068514 (2016).

66. NCBI Sequence Read Archive, https://identifiers.org/ncbi/insdc.sra:SRP068608 (2016).

67. NCBI Sequence Read Archive, https://identifiers.org/ncbi/insdc.sra:SRP068620 (2016).

68. NCBI Sequence Read Archive, https://identifiers.org/ncbi/insdc.sra:SRP068654 (2016).

69. NCBI Sequence Read Archive, https://identifiers.org/ncbi/insdc.sra:SRP069065 (2017).

70. NCBI Sequence Read Archive, https://identifiers.org/ncbi/insdc.sra:SRP069742 (2017).

71. NCBI Sequence Read Archive, https://identifiers.org/ncbi/insdc.sra:SRP070568 (2016).

72. NCBI Sequence Read Archive, https://identifiers.org/ncbi/insdc.sra:SRP073070 (2016).

73. NCBI Sequence Read Archive, https://identifiers.org/ncbi/insdc.sra:SRP073265 (2017).

74. NCBI Sequence Read Archive, https://identifiers.org/ncbi/insdc.sra:SRP074055 (2016).

75. NCBI Sequence Read Archive, https://identifiers.org/ncbi/insdc.sra:SRP074496 (2016).

76. NCBI Sequence Read Archive, https://identifiers.org/ncbi/insdc.sra:SRP075989 (2017).

77. NCBI Sequence Read Archive, https://identifiers.org/ncbi/insdc.sra:SRP079403 (2017).

78. NCBI Sequence Read Archive, https://identifiers.org/ncbi/insdc.sra:SRP079521 (2018).

79. NCBI Sequence Read Archive, https://identifiers.org/ncbi/insdc.sra:SRP080210 (2016).

80. NCBI Sequence Read Archive, https://identifiers.org/ncbi/insdc.sra:SRP080428 (2017).

81. NCBI Sequence Read Archive, https://identifiers.org/ncbi/insdc.sra:SRP080680 (2017).

82. NCBI Sequence Read Archive, https://identifiers.org/ncbi/insdc.sra:SRP082472 (2017).

83. NCBI Sequence Read Archive, https://identifiers.org/ncbi/insdc.sra:SRP082976 (2017).

84. NCBI Sequence Read Archive, https://identifiers.org/ncbi/insdc.sra:SRP083394 (2017).

85. NCBI Sequence Read Archive, https://identifiers.org/ncbi/insdc.sra:SRP083434 (2017).

86. NCBI Sequence Read Archive, https://identifiers.org/ncbi/insdc.sra:SRP083901 (2017).

87. NCBI Sequence Read Archive, https://identifiers.org/ncbi/insdc.sra:SRP087715 (2017).

88. NCBI Sequence Read Archive, https://identifiers.org/ncbi/insdc.sra:SRP090261 (2016).

89. NCBI Sequence Read Archive, https://identifiers.org/ncbi/insdc.sra:SRP090335 (2017).

90. NCBI Sequence Read Archive, https://identifiers.org/ncbi/insdc.sra:SRP090490 (2017).

91. NCBI Sequence Read Archive, https://identifiers.org/ncbi/insdc.sra:SRP090651 (2017).

92. NCBI Sequence Read Archive, https://identifiers.org/ncbi/insdc.sra:SRP091741 (2017).

93. NCBI Sequence Read Archive, https://identifiers.org/ncbi/insdc.sra:SRP091855 (2018).

94. NCBI Sequence Read Archive, https://identifiers.org/ncbi/insdc.sra:SRP091867 (2016).

95. NCBI Sequence Read Archive, https://identifiers.org/ncbi/insdc.sra:SRP092609 (2019).

96. NCBI Sequence Read Archive, https://identifiers.org/ncbi/insdc.sra:SRP092777 (2017).

97. NCBI Sequence Read Archive, https://identifiers.org/ncbi/insdc.sra:SRP093592 (2017).

98. NCBI Sequence Read Archive, https://identifiers.org/ncbi/insdc.sra:SRP093928 (2017).

99. NCBI Sequence Read Archive, https://identifiers.org/ncbi/insdc.sra:SRP094708 (2017).

100. NCBI Sequence Read Archive, https://identifiers.org/ncbi/insdc.sra:SRP097883 (2017).

101. NCBI Sequence Read Archive, https://identifiers.org/ncbi/insdc.sra:SRP101553 (2017).

102. NCBI Sequence Read Archive, https://identifiers.org/ncbi/insdc.sra:SRP101605 (2018).

103. NCBI Sequence Read Archive, https://identifiers.org/ncbi/insdc.sra:SRP102378 (2017).

104. NCBI Sequence Read Archive, https://identifiers.org/ncbi/insdc.sra:SRP102417 (2018).

105. NCBI Sequence Read Archive, https://identifiers.org/ncbi/insdc.sra:SRP102775 (2018).

106. NCBI Sequence Read Archive, https://identifiers.org/ncbi/insdc.sra:SRP106137 (2018).

107. NCBI Sequence Read Archive, https://identifiers.org/ncbi/insdc.sra:SRP106774 (2018).

108. NCBI Sequence Read Archive, https://identifiers.org/ncbi/insdc.sra:SRP107174 (2017).

109. NCBI Sequence Read Archive, https://identifiers.org/ncbi/insdc.sra:SRP107743 (2017).

110. NCBI Sequence Read Archive, https://identifiers.org/ncbi/insdc.sra:SRP109164 (2017).

111. NCBI Sequence Read Archive, https://identifiers.org/ncbi/insdc.sra:SRP109773 2017).

112. NCBI Sequence Read Archive, https://identifiers.org/ncbi/insdc.sra:SRP110522 (2017).

113. NCBI Sequence Read Archive, https://identifiers.org/ncbi/insdc.sra:SRP110810 (2017).

114. NCBI Sequence Read Archive, https://identifiers.org/ncbi/insdc.sra:SRP113348 (2018).

115. NCBI Sequence Read Archive, https://identifiers.org/ncbi/insdc.sra:SRP114697 (2017).

116. NCBI Sequence Read Archive, https://identifiers.org/ncbi/insdc.sra:SRP114821 (2018).

117. NCBI Sequence Read Archive, https://identifiers.org/ncbi/insdc.sra:SRP115350 (2018).

118. NCBI Sequence Read Archive, https://identifiers.org/ncbi/insdc.sra:SRP115464 (2018).

119. NCBI Sequence Read Archive, https://identifiers.org/ncbi/insdc.sra:SRP115599 (2018).

120. NCBI Sequence Read Archive, https://identifiers.org/ncbi/insdc.sra:SRP117302 (2018).

121. NCBI Sequence Read Archive, https://identifiers.org/ncbi/insdc.sra:SRP118875 (2018).

122. NCBI Sequence Read Archive, https://identifiers.org/ncbi/insdc.sra:SRP118960 (2018).

123. NCBI Sequence Read Archive, https://identifiers.org/ncbi/insdc.sra:SRP119174 (2017).

124. NCBI Sequence Read Archive, https://identifiers.org/ncbi/insdc.sra:SRP125864 (2016).

125. NCBI Sequence Read Archive, https://identifiers.org/ncbi/insdc.sra:SRP132277 (2018).

126. NCBI Sequence Read Archive, https://identifiers.org/ncbi/insdc.sra:SRP132591 (2018).

127. NCBI Sequence Read Archive, https://identifiers.org/ncbi/insdc.sra:SRP132598 (2018).

128. NCBI Sequence Read Archive, https://identifiers.org/ncbi/insdc.sra:SRP136886 (2012).

129. NCBI Sequence Read Archive, https://identifiers.org/ncbi/insdc.sra:SRP139483 (2019).

130. NCBI Sequence Read Archive, https://identifiers.org/ncbi/insdc.sra:SRP142723 (2018).

131. NCBI Sequence Read Archive, https://identifiers.org/ncbi/insdc.sra:SRP148813 (2018).

132. NCBI Sequence Read Archive, https://identifiers.org/ncbi/insdc.sra:SRP150527 (2019).

133. NCBI Sequence Read Archive, https://identifiers.org/ncbi/insdc.sra:SRP151262 (2018).

134. NCBI Sequence Read Archive, https://identifiers.org/ncbi/insdc.sra:SRP153934 (2018).

135. NCBI Sequence Read Archive, https://identifiers.org/ncbi/insdc.sra:SRP160913 (2018).

136. NCBI Sequence Read Archive, https://identifiers.org/ncbi/insdc.sra:SRP161632 (2018).

137. NCBI Sequence Read Archive, https://identifiers.org/ncbi/insdc.sra:SRP195764 (2019).

138. European Nucleotide Archive, https://identifiers.org/ena.embl:ERP001713 (2014).

139. European Nucleotide Archive, https://identifiers.org/ena.embl:ERP003251 (2013).

140. European Nucleotide Archive, https://identifiers.org/ena.embl:ERP003790 (2015).

141. European Nucleotide Archive, https://identifiers.org/ena.embl:ERP005177 (2015).

142. European Nucleotide Archive, https://identifiers.org/ena.embl:ERP005905 (2015).

143. European Nucleotide Archive, https://identifiers.org/ena.embl:ERP009341 (2015).

144. European Nucleotide Archive, https://identifiers.org/ena.embl:ERP010027 (2017).

145. European Nucleotide Archive, https://identifiers.org/ena.embl:ERP010084 (2016).

146. European Nucleotide Archive, https://identifiers.org/ena.embl:ERP010743 (2016).

147. European Nucleotide Archive, https://identifiers.org/ena.embl:ERP011924 (2016).

148. European Nucleotide Archive, https://identifiers.org/ena.embl:ERP012017 (2016).

149. European Nucleotide Archive, https://identifiers.org/ena.embl:ERP013208 (2016).

150. European Nucleotide Archive, https://identifiers.org/ena.embl:ERP013987 (2017).

151. European Nucleotide Archive, https://identifiers.org/ena.embl:ERP014227 (2016).

152. European Nucleotide Archive, https://identifiers.org/ena.embl:ERP017480 (2018).

153. European Nucleotide Archive, https://identifiers.org/ena.embl:ERP017851 (2017).

154. European Nucleotide Archive, https://identifiers.org/ena.embl:ERP017915 (2017).

155. European Nucleotide Archive, https://identifiers.org/ena.embl:ERP019580 (2017).

156. European Nucleotide Archive, https://identifiers.org/ena.embl:ERP019924 (2017).

157. European Nucleotide Archive, https://identifiers.org/ena.embl:ERP020657 (2017).

158. European Nucleotide Archive, https://identifiers.org/ena.embl:ERP022511 (2019).

159. European Nucleotide Archive, https://identifiers.org/ena.embl:ERP022742 (2017).

160. European Nucleotide Archive, https://identifiers.org/ena.embl:ERP023275 (2018).

161. European Nucleotide Archive, https://identifiers.org/ena.embl:ERP023718 (2018).

162. European Nucleotide Archive, https://identifiers.org/ena.embl:ERP023855 (2018).

163. European Nucleotide Archive, https://identifiers.org/ena.embl:ERP106131 (2018).

164. European Nucleotide Archive, https://identifiers.org/ena.embl:ERP107634 (2019).

165. European Nucleotide Archive, https://identifiers.org/ena.embl:ERP107636 (2019).

166. European Nucleotide Archive, https://identifiers.org/ena.embl:ERP110188 (2019).

167. European Nucleotide Archive, https://identifiers.org/ena.embl:ERP112007 (2019).

168. DNA Data Bank of Japan, https://trace.ddbj.nig.ac.jp/DRASearch/submission?acc=DRA000926 (2014).

169. DNA Data Bank of Japan, https://trace.ddbj.nig.ac.jp/DRASearch/submission?acc=DRA000937 (2014).

170. DNA Data Bank of Japan, https://trace.ddbj.nig.ac.jp/DRASearch/submission?acc=DRA001737 (2016).

171. DNA Data Bank of Japan, https://trace.ddbj.nig.ac.jp/DRASearch/submission?acc=DRA002424 (2016).

172. DNA Data Bank of Japan, https://trace.ddbj.nig.ac.jp/DRASearch/submission?acc=DRA002469 (2016).

173. DNA Data Bank of Japan, https://trace.ddbj.nig.ac.jp/DRASearch/submission?acc=DRA003024 (2016).

174. DNA Data Bank of Japan, https://trace.ddbj.nig.ac.jp/DRASearch/submission?acc=DRA003730 (2016).

175. DNA Data Bank of Japan, https://trace.ddbj.nig.ac.jp/DRASearch/submission?acc=DRA004913 (2017).

176. DNA Data Bank of Japan, https://trace.ddbj.nig.ac.jp/DRASearch/submission?acc=DRA006519 (2018).

177. DNA Data Bank of Japan, https://trace.ddbj.nig.ac.jp/DRASearch/study?acc=DRP002783 (2015).

178. DNA Data Bank of Japan, https://trace.ddbj.nig.ac.jp/DRASearch/study?acc=DRP003138 (2016).

179. DNA Data Bank of Japan, https://trace.ddbj.nig.ac.jp/DRASearch/study?acc=DRP005365 (2019).

180. Dryad Digital Repository, https://doi.org/10.5061/dryad.2fc32 (2015).

181. Dryad Digital Repository, http://dx.doi.org/10.5061/dryad.n82g9 (2017).

182. Dryad Digital Repository, http://dx.doi.org/10.5061/dryad.2343k (2015).

183. Dryad Digital Repository, https://doi.org/10.5061/dryad.gp302 (2015).

184. Dryad Digital Repository, http://dx.doi.org/10.5061/dryad.cq2rb (2016).

185. Dryad Digital Repository, https://doi.org/10.5061/dryad.8fn8j (2017).

186. Dryad Digital Repository, https://doi.org/10.5061/dryad.216tp (2013).

187. MG-RAST, https://www.mg-rast.org/mgmain.html?mgpage=search&search=4783710.3 (2012).

188. MG-RAST, https://www.mg-rast.org/mgmain.html?mgpage=search&search=4702703.3 (2016).

189. MG-RAST, https://www.mg-rast.org/mgmain.html?mgpage=search&search=4524551.3 (2014).

190. MG-RAST, https://www.mg-rast.org/mgmain.html?mgpage=search&search=4544233.3 (2016).

191. MG-RAST, https://www.mg-rast.org/mgmain.html?mgpage=search&search=4683808.3 (2017).

192. MG-RAST, https://www.mg-rast.org/mgmain.html?mgpage=search&search=4684008.3 (2016).

193. MG-RAST, https://www.mg-rast.org/mgmain.html?mgpage=search&search=4696490.3 (2016).

194. MG-RAST, https://www.mg-rast.org/mgmain.html?mgpage=search&search=4715213.3 (2016).

195. MG-RAST, https://www.mg-rast.org/mgmain.html?mgpage=search&search=4563787.3 (2015).

196. MG-RAST, https://www.mg-rast.org/mgmain.html?mgpage=search&search=4563788.3 (2015).

197. MG-RAST, https://www.mg-rast.org/mgmain.html?mgpage=search&search=4620497.3 (2015).

198. MG-RAST, https://www.mg-rast.org/mgmain.html?mgpage=search&search=4620498.3 (2015).

199. MG-RAST, https://www.mg-rast.org/mgmain.html?mgpage=search&search=mgp13293 (2012).

200. MG-RAST, https://www.mg-rast.org/mgmain.html?mgpage=search&search=mgp1617 (2013).

201. MG-RAST, https://www.mg-rast.org/mgmain.html?mgpage=search&search=mgp9003 (2014).

202. PlutoF repository, https://plutof.ut.ee/#/filerepository/view/1561672 (2017).

203. PlutoF repository, https://plutof.ut.ee/#/filerepository/view/1562683 (2017).

204. PlutoF repository, https://plutof.ut.ee/#/doi/10.15156/BIO/100002 (2015).

205. PlutoF repository, https://plutof.ut.ee/#/doi/10.15156/BIO/587446 (2017).

206. UNITE database, http://unite.ut.ee/454_EcM_CMR.zip (2010).

207. Schappe, T., Albornoz, F.E., Turner, B.L., Neat, A., Condit, R. and Jones, F.A. TLS: uncultured fungus internal transcribed spacer 1, targeted locus study. Genbank https://identifiers.org/ncbi/insdc:KAYV00000000.1 (2017).

208. Schappe, T., Albornoz, F.E., Turner, B.L., Neat, A., Condit, R. and Jones, F.A. TLS: uncultured fungus internal transcribed spacer 1, targeted locus study. Genbank https://identifiers.org/ncbi/insdc:KAYU00000000.1 (2017).

209. Schappe, T., Albornoz, F.E., Turner, B.L., Neat, A., Condit, R. and Jones, F.A. TLS: uncultured fungus internal transcribed spacer 1, targeted locus study. Genbank https://identifiers.org/ncbi/insdc:KAYT00000000.1 (2017).

210. Vaz, A.B., Fonseca, P.L., Leite, L.R., Badotti, F., Salim, A.C., Araujo, F.M., Cuadros-Orellana, S., Duarte, A.A., Rosa, C.A., Oliveira, G. and Goes-Neto, A. MIMS Environmental/Metagenome sample from biofilm metagenome. Genbank https://identifiers.org/ncbi/insdc:SAMN02934078 (2017).

211. Vaz, A.B., Fonseca, P.L., Leite, L.R., Badotti, F., Salim, A.C., Araujo, F.M., Cuadros-Orellana, S., Duarte, A.A., Rosa, C.A., Oliveira, G. and Goes-Neto, A. MIMS Environmental/Metagenome sample from biofilm metagenome. Genbank https://identifiers.org/ncbi/insdc:SAMN02934079 (2017).

212. Australian Antarctic Data Center, http://dx.doi.org/10.4225/15/526f42ada05b1 (2016).

213. Hartmann, M., Howes, C.G., VanInsberghe, D., Yu, H., Bachar, D., Christen, R., Nilsson, R.H., Hallam, S.J. and Mohn, W.W. Significant and persistent impact of timber harvesting on soil microbial communities in Northern coniferous forests. Supplementary Data 2 https://static-content.springer.com/esm/art%3A10.1038%2Fismej.2012.84/MediaObjects/41396_2012_BFismej201284_MOESM78_ESM.zip (2012).

214. Rime, T., Hartmann, M. and Frey, B. Potential sources of microbial colonizers in an initial soil ecosystem after retreat of an alpine glacier. Supplementary Information https://static-content.springer.com/esm/art%3A10.1038%2Fismej.2015.238/MediaObjects/41396_2016_BFismej2015238_MOESM67_ESM.zip (2016).

215. Czech Academy of Sciences, www.biomed.cas.cz/mbu/lbwrf/metastudy_datasets/GF0001.zip.

216. Czech Academy of Sciences, www.biomed.cas.cz/mbu/lbwrf/metastudy_datasets/GF0002.zip,

217. Czech Academy of Sciences, www.biomed.cas.cz/mbu/lbwrf/metastudy_datasets/GF0003.zip,

218. Czech Academy of Sciences, www.biomed.cas.cz/mbu/lbwrf/metastudy_datasets/GF0004.zip,

219. Czech Academy of Sciences, www.biomed.cas.cz/mbu/lbwrf/metastudy_datasets/GF0005.zip,

220. Czech Academy of Sciences, www.biomed.cas.cz/mbu/lbwrf/metastudy_datasets/GF0006.zip,

221. Czech Academy of Sciences, www.biomed.cas.cz/mbu/lbwrf/metastudy_datasets/GF0007.zip,

222. Czech Academy of Sciences, www.biomed.cas.cz/mbu/lbwrf/metastudy_datasets/GF0008.zip,

223. Czech Academy of Sciences, www.biomed.cas.cz/mbu/lbwrf/metastudy_datasets/GF0009.zip,

224. Czech Academy of Sciences, www.biomed.cas.cz/mbu/lbwrf/metastudy_datasets/GF0010.zip,

225. Czech Academy of Sciences, www.biomed.cas.cz/mbu/lbwrf/metastudy_datasets/GF0011.zip,

226. Czech Academy of Sciences, www.biomed.cas.cz/mbu/lbwrf/metastudy_datasets/GF0012.zip.

227. Anslan, S., Nilsson, R.H., Wurzbacher, C., Baldrian, P., Tedersoo, L. & Bahram, M. Great differences in performance and outcome of high-throughput sequencing data analysis platforms for fungal metabarcoding. Mycokeys 39, 29–40 (2018).

